# Reduced function of the adaptor SH2B3 promotes T1D via altered gc cytokine-regulated, T cell intrinsic immune tolerance

**DOI:** 10.1101/2024.08.02.606362

**Authors:** Taylor K. Watson, Aaron B.I. Rosen, Travis Drow, Jacob A. Medjo, Matthew A. MacQuivey, Yan Ge, H. Denny Liggitt, Dane A. Grosvenor, Kimberly A. Dill-McFarland, Matthew C. Altman, Patrick J. Concannon, Jane H. Buckner, David J. Rawlings, Eric J. Allenspach

## Abstract

Genome-wide association studies have identified *SH2B3* as an important non-MHC gene for islet autoimmunity and type 1 diabetes (T1D). In this study, we found a single *SH2B3* haplotype significantly associated with increased risk for human T1D, and this haplotype carries the single nucleotide variant rs3184504*T in *SH2B3.* To better characterize the role of SH2B3 in T1D, we used mouse modeling and found a T cell-intrinsic role for SH2B3 regulating peripheral tolerance. SH2B3 deficiency had minimal effect on TCR signaling or proliferation across antigen doses, yet enhanced cell survival and cytokine signaling including common gamma chain-dependent and interferon-gamma receptor signaling. SH2B3 deficient CD8+T cells showed augmented STAT5-MYC and effector-related gene expression partially reversed with blocking autocrine IL-2 in culture. Using the RIP-mOVA model, we found CD8+ T cells lacking SH2B3 promoted early islet destruction and diabetes without requiring CD4+ T cell help. SH2B3-deficient cells demonstrated increased survival post-transfer compared to control cells despite a similar proliferation profile in the same host. Next, we created a spontaneous NOD*.Sh2b3^-/-^*mouse model and found markedly increased incidence and accelerated T1D across sexes. Collectively, these studies identify SH2B3 as a critical mediator of peripheral T cell tolerance limiting the T cell response to self-antigens.

**Article Highlights:**

- The rs3184504 polymorphism, encoding a hypomorphic variant of the negative regulator SH2B3, strongly associates with T1D.
- SH2B3 deficiency results in hypersensitivity to cytokines, including IL-2, in murine CD4+ and CD8+ T cells.
- SH2B3 deficient CD8+ T cells exhibit a comparable transcriptome to wild-type CD8+ T cells at baseline, but upon antigen stimulation SH2B3 deficient cells upregulate genes characteristic of enhanced JAK/STAT signaling and effector functions.
- We found a T-cell intrinsic role of SH2B3 leading to severe islet destruction in an adoptive transfer murine T1D model, while global SH2B3 deficiency accelerated spontaneous NOD diabetes across sexes.

## INTRODUCTION

Type 1 Diabetes (T1D) is caused by T-cell mediated destruction of pancreatic β-cells resulting in life-long insulin dependence. T1D is a polygenic disease with >90% of subjects carrying a high-risk human leukocyte antigen (HLA) together with additional non-HLA risk alleles^1^. Several genetic risk variants are now being combined into a genetic risk score to prospectively identify high-risk individuals or predict T1D disease progression^1–3^. A priority credible causative variant for T1D^4^, *rs3184504*, that encodes a missense substitution in the *SH2B3* gene has also been associated with risk for other autoimmune conditions including rheumatoid arthritis^5^, celiac disease^6^, systemic lupus erythematosus^7^, and multiple sclerosis^8^, as well as inflammatory conditions like cardiovascular disease^9^. Although *SH2B3* has been associated with autoimmune diseases, the cellular mechanisms that contribute to autoimmune pathogenesis are yet to be elucidated.

SH2B adaptor protein 3 (SH2B3; also known as LNK) is a dose-dependent intracellular adaptor and negative regulator of cytokine receptor signaling^10^. SH2B3 regulates hematopoiesis by altering signaling downstream of JAK2 and JAK3-dependent receptors as well as several receptor tyrosine kinases. JAK-STAT pathways are critical for β-cell destruction in spontaneous diabetes^11^ and JAK inhibitors targeting these pathways have shown efficacy in preserving β-cell function or even reversing diabetes in the NOD mouse model^12^ and in human T1D subjects^13^. Recent work has shown that SH2B3 exerts negative regulation via control of JAK stability and degradation^10,14^. Targeted testing in *Sh2b3^-/-^* mice has shown hyperresponsiveness to common gamma-chain (γ_c_) cytokines, including IL-7 and IL-15, as well as Th1 cytokines, like IL-12 and IFN-γ^15–18^. However, the cell types and pathways impacted by SH2B3 in T1D are not clear. In this study, we tested the association of rs3184504, a coding variant in *SH2B3*, with human T1D and performed a haplotype analysis indicating that increased risk associates with a single nucleotide variant rs3184504*T in *SH2B3*. Given the variant encodes a hypomorphic SH2B3 adaptor, we next tested both global and conditional *Sh2b3* knockout mice in several murine models of T1D and in detailed in vitro studies. We identified a unique role for SH2B3 in regulating early cytokine signaling in the immediate phase post-T cell activation. Next, we demonstrate that SH2B3 modulates peripheral CD8+ T cell tolerance via altering the response to self-antigen and the competitive fitness of self-reactive T cells in vivo wherein loss of SH2B3 function promotes both increased survival and pathogenicity. Collectively, our data highlight the importance of the SH2B3 risk allele in T1D and in other autoimmune diseases and provide new mechanistic insight into a key signaling function of SH2B3.

## MATERIALS & METHODS

### Association and haplotype analysis

Genomic DNA from T1D-affected sibling pairs (ASP) and trio families were obtained from the Type 1 Diabetes Genetics Consortium (T1DGC) as described previously^19^. Association analysis utilized data from T1DGC using genotypes for the 12q24 region. The Family-Based Association Test (FBAT) program (version 2.0.4) was used for single-marker association tests and haplotype analyses. Minor allele frequencies were estimated using PLINK (v.1.90).

### Mice

All animal care and experimentation occurred with Institutional Animal Care and Use Committee approval at the SCRI Animal Facility. *Sh2b3^-/-^*and *Sh2b3^fl/fl^* mice were described previously^20^. All other strains were purchased from Jackson Laboratories (Bar Harbor, Maine): RIP-mOVA, CD4Cre, B6 CD45.1, Ai14, OT-I, OT-II, and NOD. The NOD.*Sh2b3^-/^*^-^ mice were generated by backcrossing *Sh2b3^-/-^* mice 15 generations to NOD with genome wide SNP validation including probes flanking the *Sh2b3* locus. NOD.*Sh2b3^+/-^*mice were intercrossed to create littermate cohorts of each gender and genotype experimentally aged up to 40 weeks. Mice were otherwise used at 6-12 weeks of age.

### Flow cytometry

Cells were stained as indicated for surface and intracellular markers, viability or proliferative dyes per manufacturer protocols; cells were fixed via 2% PFA and permeabilized with Perm Buffer III (details in **Supplementary Material**). Fluorescently labeled cells were acquired on BD LSR II or Fortessa (Becton Dickson) and analyzed using FlowJo (v10.8.1) (Treestar, Ashland, OR). Graphs and figures were prepared using GraphPad Prism v10.0.0 (Boston, MA) and Adobe Illustrator v16.0.0, respectively.

### Cell cultures

Spleen and/or lymph node (LN) cells were purified by negative selection for total or naïve CD8+ or CD4+ T cells (Stem Cell Technologies). Bulk splenocytes or purified naïve T cells were plated with stimulatory anti-CD3ε and anti-CD28 antibodies with or without anti–IL-2 blocking antibody at indicated doses. Naïve CD8+ T cells were co-cultured with irradiated (3500rad) splenocytes pulsed with SIINFEKL (N4) or SIIQFEKL (Q4) (AnaSpec). Splenocytes were rested for 2 hours in serum-free RPMI prior to cytokine stimulationwith recombinant murine IL-2, IL-15, IFN-γ (Peprotech) and in-house IL-7^21^ at indicated doses.

### Diabetes murine studies

Purified naïve CD8+ OT-I cells from spleen and cutaneous LN were transferred retro-orbitally into sex-matched RIP-mOVA^+^ hosts (7-14 weeks) with co-transfer (1:1) or separate transfers. Tissues were fixed in 10% neutral buffered formalin prior to paraffin-embedding and sectioning with indicated staining and immunohistochemistry. Pancreases were digested in collagenase IV (2 mg/mL) and PBMCs were purified via percoll density gradient prior to flow cytometry. Diabetes cohorts were monitored for hyperglycemia characterized by blood glucose >300 mg/dL measured, and two consecutive readings required euthanasia per protocol.

### Expression analysis

SH2B3 mRNA (ENSG00000111252) transcript sequence analysis was described previously^22,23^. Murine *Sh2b3* and *Actb* transcripts were assessed by RT-PCR analysis using SYBR green per manufacturer protocol (Thermo). Bulk mRNA-seq was performed via NovaSeq PE150 RNA sequencing and analyzed via Biojupies^24^ or separately as described^25^ (details in **Supplementary Material**).

### Statistical analysis

All statistical analysis used GraphPad Prism version 7.0b except where noted. All specific statistical tests and *P*-values are indicated in the relevant figures.

## RESULTS

### The human SH2B3^R262W^ variant associates with T1D as a single haplotype

Initial reports on the association of the 12q24.12 region, containing *SH2B3*, with T1D were unable to unambiguously identify the causative variant in the region from genetic data^26^. Two variants in the region have been credibly associated with autoimmunity^4^, namely rs3184504*C>T (**<u>C</u>**GG>**<u>T</u>**GG) encoding SH2B3^R262W^ and rs653178*G>A in an untranslated region of *ATXN2*. Subsequent studies utilizing an expanded number of subjects with more diverse genetic backgrounds have prioritized rs3184504 as the most credible causative variant for in this region^4^. Separately, we confirmed the association of this locus with T1D (**Supplementary Table I-II**). Our family-based haplotype association analysis identified two T1D-associated haplotypes in this cohort, H1 and H2. The H1 haplotype carrying the T allele of rs3184504 and the G allele of rs653178 conferred risk for T1D. SH2B3^R262W^ is the major variant in most European populations and is protective in sepsis^20^, thus is predicted to be enriched due to positive selection. Given linkage disequilibrium in the region, enrichment is also observed for rs653178*G^26–28^. Thus, a single SH2B3^RISK^ haplotype strongly associated with T1D in our cohort.

### SH2B3 expression is rapidly induced in T cells following antigen stimulation

Our previous work demonstrated that SH2B3 exhibits dose dependent adaptor activity^20,30^. Previous work has also shown that *SH2B3* expression can be induced in response to cytokine signaling^31^. While *SH2B3* is expressed at high levels in peripheral blood monocytes, mature blood B and T lymphocytes exhibit minimal expression^22,32^. Thus, we hypothesized that stimulation was required to mediate increased *SH2B3* expression in lymphocytes. To test this, we analyzed public datasets for *SH2B3* expression following stimulation. Human naïve CD4+ and CD8+ T cells^22^, as well as memory CD4+ cells^23^, upregulated *SH2B3* transcripts within 2-4 hours of co-stimulation (**Fig. 1A-B**). Using a murine model system, we validated that TCR stimulation alone was sufficient to rapidly upregulate *Sh2b3* expression, with or without blocking autocrine IL-2 (**Fig. 1C**). Additionally, IL-2 alone was sufficient to induce *Sh2b3* expression. Western blot analysis showed SH2B3 protein upregulation at 6-8 hours following antigen stimulation **(Supplementary Fig. 1)**. Thus, SH2B3 is an inducible adaptor rapidly expressed in T cells.

**Figure 1.**
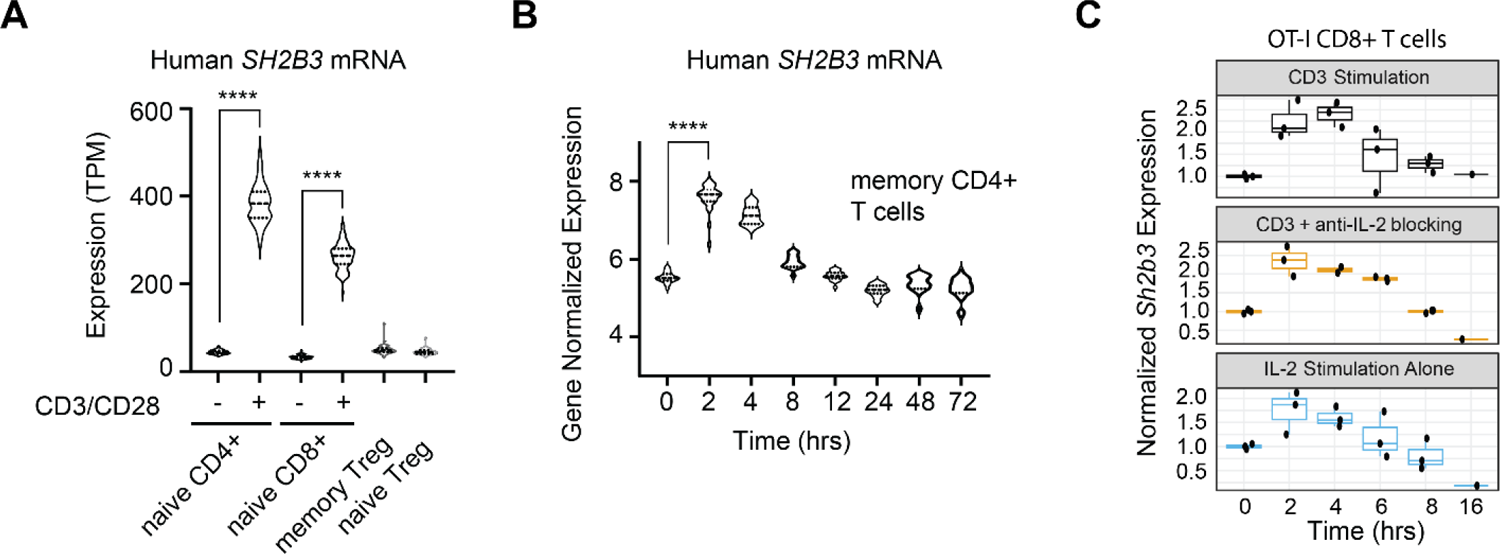
SH2B3 is rapidly upregulated after stimulation in both human and mouse T cells. (**A** - **B**) Focused *SH2B3* expression analysis of public mRNA-seq data from (A) Naïve CD4+ T cells (CD3^+^CD4^+^CD45RA^+^CCR7^+^), naïve CD8+ T cells (CD3^+^CD8a^+^CD45RA^+^CCR7^+^), naïve Treg (CD3^+^CD4^+^CD45RA^+^CD25^+^CD127^-^) or memory Treg (CD3^+^CD4^+^CD45RA^-^CD25^+^ CD127-) cells were purified from human PBMC (n=88 donors) as published^22,23^. Naïve T cells were unstimulated or stimulated using Human T-Activator anti-CD3/anti-CD28 beads at a ratio of 1:1 for 4 hours. (B) RNA-seq time series experiment on CD4+ memory T cells activated with anti-CD3/CD28 beads in individuals of European ancestry with no autoimmune disease (n=24) as published^23^. Cells were stimulated with anti-CD3/CD28 beads and collected at 0, 2, 4, 8, 12, 24, 48, and 72 hours. (**C**) Purified mouse naïve CD8+ T cells from OT-I.*Sh2b3^+/+^* mice were stimulated for 0, 2, 4, 6, 8, and 16 hours with anti-CD3 (10ug/mL) +/- blocking anti-IL-2 (10ug/mL) or IL-2 (100ng/mL) prior to mRNA isolation and RT-PCR for *Sh2b3/Actb* expression (n=3/genotype; representative of n=3 experiments).

### SH2B3 deficient mice exhibit normal thymic and mildly altered peripheral T cell development

SH2B3 was originally described to regulate TCR signaling^33^; however, subsequent studies found *Sh2b3^-/-^* mice had normal baseline thymocyte development and peripheral T cell numbers^34^. Similarly, we found no significant changes in thymocyte populations in either global (*Sh2b3^-/-^*) or lineage-specific (CD4-Cre x *Sh2b3^flox/flox^*) mice compared to littermate controls **(Supplementary Fig. 2A-B)**. T cell development was also comparable across genotypes when crossed to OT-I or OT-II TCR transgenic backgrounds, suggesting that SH2B3 is not necessary for proximal TCR signaling **(Supplementary Fig. 2C-D)**. In the periphery, *Sh2b3^-/-^* mice exhibited modest expansion of CD44^+^CD62L^+^CD8^+^ T cells. This phenotype was also present in T cell lineage-specific knockout mice where the majority of these CD8+ T cells were CD49d^lo^, consistent with virtual or innate-like memory CD8+ T cells (**Supplementary Fig.2**). The proportion of regulatory T cells (Treg) were significantly increased in *Sh2b3^-/-^* mice compared to controls, suggesting enhanced IL-2 signaling **(Supplementary Fig. 2E–I)**. In summary, while SH2B3 was not necessary for thymocyte development or generation of peripheral naïve T cell subsets, SH2B3-deficient mice exhibited a modest expansion of T lineage populations known to be preferentially responsive to γ_c_ cytokines.

### SH2B3 negatively regulates γ_c_ cytokine responses in primary mouse T cells

Next, we directly compared the responsiveness of naïve and memory splenic CD4+ and CD8+ T cells from Sh2b3^+/+^ vs. Sh2b3^-/-^ mice to γ_c_ cytokines. Baseline cytokine receptor surface expression (for IL-2Rα, IL-2Rβ, IL-7Rα, IL-2Rγ), total STAT5 levels, and phosphorylated (p)STAT5 and pSTAT1 levels were comparable across genotypes **(Supplementary Fig. 3A-B)**. Furthermore, we assessed the impact of stimulation with saturating doses of IL-2, IL-7, or IL-15 for up to 120 minutes (**Fig. 2**). We compared the area under the curve (AUC) for pSTAT5 levels in CD4+ and CD8+ T cell subsets from each genotype. Memory cells responded most potently to high-dose IL-2 and IL-15, while naïve T cells responded dominantly to IL-7 regardless of genotype (**Fig 2A-C**). The pSTAT5 AUC was increased in *Sh2b3^-/-^* vs. control T cells in response to all γ_c_ cytokines. The impact of SH2B3 deficiency was most evident within the naïve CD4+ and CD8+ T cell compartments (**Fig. 2A-C**).

**Figure 2.**
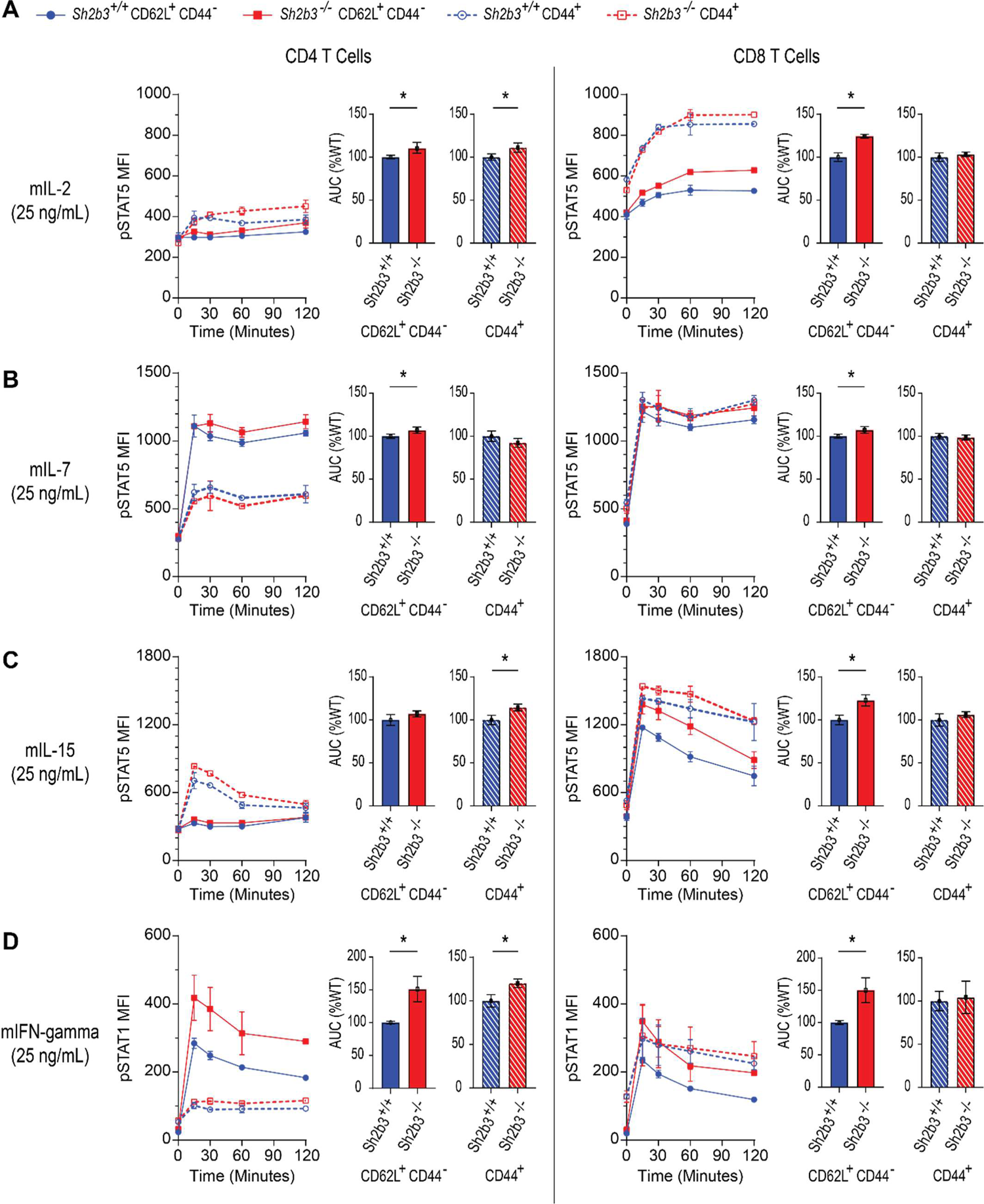
Loss of SH2B3 results in enhanced responsiveness to γ_c_ family cytokines and IFN-γ in CD4+ and CD8+ T cells. (**A** - **C**) Plots depict STAT5 and (**D**) STAT1 phosphorylation based on T cell subset following stimulation of bulk splenocytes from Sh2b3^+/+^ and Sh2b3^-/-^ mice with 25 ng/mL of (A) murine IL-2, (B) murine IL-7, (C) murine IL-15, and (D) murine IFN-γ. Bulk splenocytes were plated at 1e6/well in U-bottomed 96-well plates. Cells were stimulated for 0, 15, 30, 60, and 120 minutes at 37C. AUC was calculated in GraphPad Prism (*n* = 2, where *n* reflects the number of individual mice per genotype). Marked significance in AUC reflects 95% confidence intervals that do not overlap between Sh2b3^+/+^ and Sh2b3^-/-^ cells.

SH2B3 also regulates JAK2-regulated cytokine signaling^10^. Therefore, we stimulated T cell subsets from each genotype with IFN-γ. *Sh2b3^-/-^* T cells exhibited augmented pSTAT1 levels across T cell subsets compared to control cells (**Fig. 2D**). While baseline total STAT1 levels were increased in naïve *Sh2b3^-/-^*CD8+ T cells compared to control cells, pSTAT1 was not different across all subsets and genotypes at baseline **(Supplementary Fig. 3B)**. Collectively, these findings demonstrate that loss of SH2B3 augments T cell responsiveness to both γ_c_ family and IFN-γ cytokine signaling, even in the absence of antigen signaling, with this impact most evident in naïve T cells.

### SH2B3 regulates IL-2 responsiveness in recently activated T cells

Cytokines can impact TCR responsiveness by altering activation thresholds, differentiation, and/or effector function^35^. We next utilized CD8+ and CD4+ T cells with defined antigen receptor specificity isolated from OT-I and OT-II TCR transgenic mice, respectively, to compare responses across *Sh2b3* genotypes. First, we stimulated OT-I T cells with plate bound anti-CD3 antibodies +/- CD28 co-stimulation (**Fig. 3A-B**). In comparison to control T cells, *Sh2b3^-/-^* CD8+ T cells exhibited augmented CD25 (IL-2Rα) expression across culture conditions throughout a 2-day period. In contrast, CD69 surface expression as a proximal readout of TCR signaling did not differ across genotypes. A similar increase in CD25 expression (and lack of differential CD69 expression) was observed in *Sh2b3^-/-^* OT-II cells. The difference in CD25 expression was consistent across a dose curve of stimulation for both CD4+ and CD8+ T cells (**Fig. 3A-B; Supplementary Fig. 4A-B)**. Given that CD25 expression on T cells can be induced by either antigen stimulation or by IL-2 itself via a positive feedback loop^35^ we hypothesized that the augmented CD25 expression in *Sh2b3^-/-^* T cells may be secondary to enhanced IL-2R autocrine signaling. When adding an IL-2 blocking antibody in culture, the difference in CD25 expression between genotypes normalized (**Fig. 3A**).

**Figure 3.**
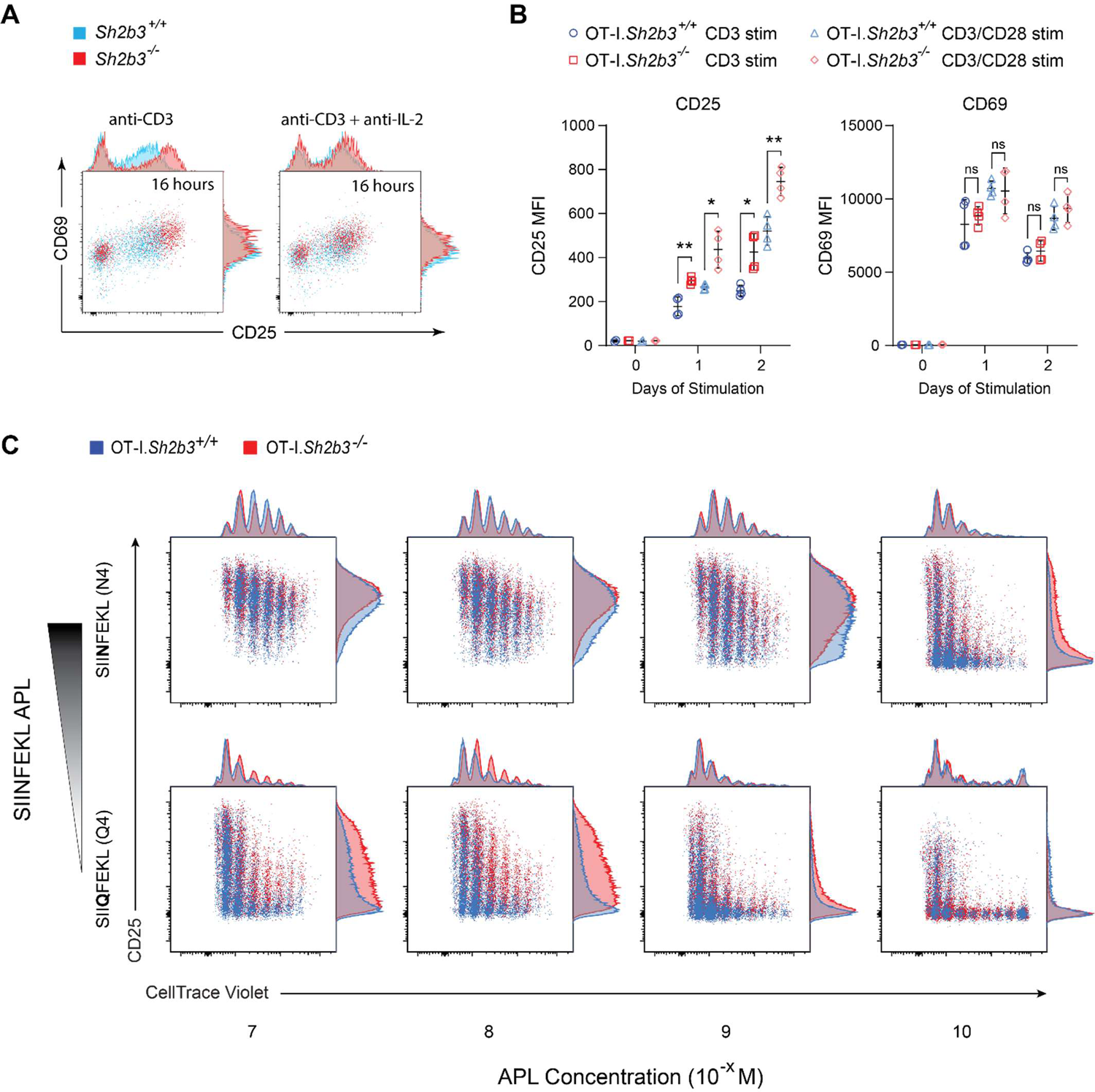
SH2B3 negatively modulates IL2R signaling while minimally impacting TCR signaling upon antigen engagement. (**A**) CD8+ T cells from OT-I.Sh2b3^+/+^ and OT-I.Sh2b3^-/-^ were stimulated with plate bound anti-CD3 stimulatory antibody (10 ug/mL) with or without anti-IL-2 blocking antibody (10 ug/mL). Plots depict CD25 (y-axis) vs. CD69 (x-axis) expression in OT-I CD8+ T cells at 16 hr post-stimulation. (**B**) Graphs depict CD25 and CD69 MFI in OT-I^+^ CD8+ T cells across a TCR stimulation timecourse. Naïve OT-I.Sh2b3^+/+^ and OT-I.Sh2b3^-/-^ CD8+ T cells were plated at 1e5/well in flat-bottomed 96-well plates coated with 3.2 ug/mL anti-CD3 stimulatory antibody with and without 3.2 ug/mL anti-CD28 stimulatory antibody for 0, 1, and 2 days. (**C**) Representative flow cytometry plots of CD25 and CellTrace Violet in OT-I.Sh2b3^+/+^ and OT-I.Sh2b3^-/-^ at varying TCR stimulation strengths. CellTrace Violet stained naïve CD8+ T cells were plated at 7.5e4/well in U-bottomed 96-well plates with irradiated APCs (1e5/well) loaded with SIINFEKL altered peptide ligands (APLs) at 10^-10^ – 10^-7^ M for 72 hrs. (B) Significance was calculated using two-tailed t tests with correction for multiple hypotheses using GraphPad Prism software (*n* = 4, where *n* reflects technical duplicates of two individual mice per genotype). *P < 0.05, **P < 0.01.

Using the alternative peptide ligands specific for OT-I T cells, we tested the function of SH2B3 across varying degrees of TCR engagement. Naïve OT-I cells from each genotype were stimulated with antigen presenting cells (APCs) pulsed with either high-affinity OVA peptide (N4) or the lower affinity peptide (Q4). CD25 expression was augmented in *Sh2b3^-/-^* vs wildtype OT-I T cells; this difference was most pronounced for T cells activated using lower affinity peptide (**Fig. 3C**). In contrast, proliferation was comparable across the dose range for both peptide ligands. Additionally, we observed no difference in secreted IL-2 across genotypes following cell activation **(Supplementary Fig. 4C-D).** These observations support that SH2B3 functions to downregulate early phase T cell activation in response to TCR engagement by negatively modulating an immediate post-activation IL-2-mediated positive feedback signaling loop rather than by altering early TCR signaling.

### SH2B3 deficient cells show increased STAT5 and MYC signatures post-activation

Antigen-stimulated CD8+ T cells exhibit heterogenous levels of CD25 that correlate with differentiation and effector function^36^. We compared the transcriptional signature in naïve OT-I.*Sh2b3^+/+^* or OT-I.*Sh2b3^-/-^* T cells at baseline and post-stimulation using bulk mRNA-seq (**Fig. 4A**). First, we compared genotypes at 0, 2, and 6 hours following anti-CD3/CD28 stimulation (**Fig. 4B; Supplementary Fig. 5A-D)**. At baseline, these populations exhibited relatively few differentially expressed genes (DEG) including *Sh2b3* transcripts and a subtle increase in effector genes (*Cxcr3, Tlr7, Ifit3b*). Consistent with the lack of impact on CD69 expression, a pattern of comparable immediate early gene expression (including genes previously described in the OT-I model^37^) was observed between *Sh2b3* genotypes. Expression of the lactate transporter, *Slc15a2*, was the only significant difference across both early time points (**Supplementary Fig. 5B-C)**.

**Figure 4.**
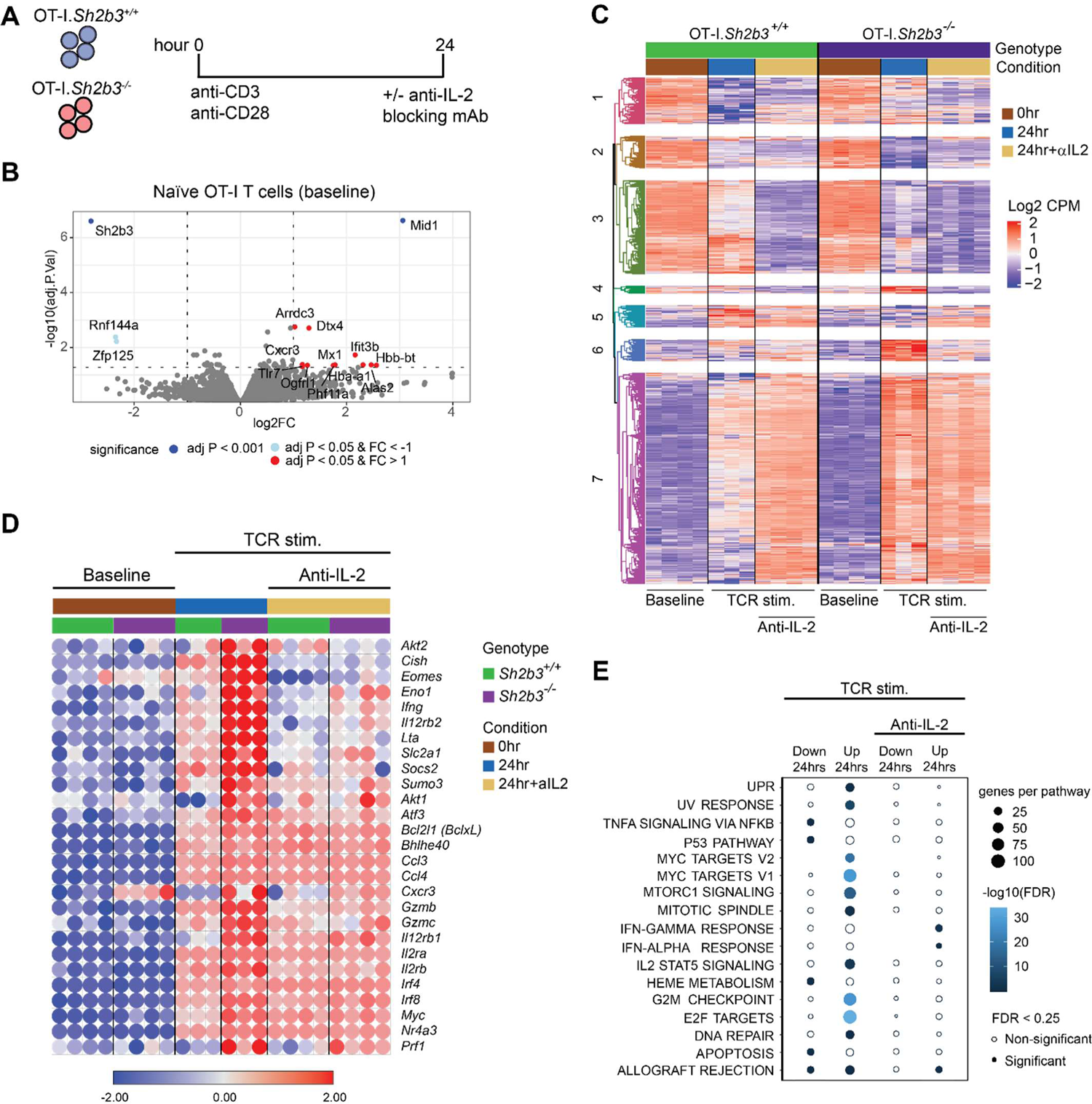
Sh2b3-deficient T cells exhibit augmented STAT5 signaling despite negligible baseline transcriptional differences. (**A**) Naïve CD8+ T cells were isolated from OT-I.Sh2b3^+/+^ and OT-I.Sh2b3^-/-^ mice and plated with anti-CD3 (10 ug/mL) and anti-CD28 (5 ug/mL) stimulatory antibodies for 0, 2, 6, and 24 hours with and without anti-IL-2 blocking antibody (10ug/mL) before RNA was isolated and sequenced. (**B**) Volcano plot depicts differentially expressed genes at baseline between OT-I.Sh2b3^+/+^ and OT-I.Sh2b3^-/-^ CD8+ T cells. (**C** – **E**) Bulk RNAseq data on expression in naïve CD8+ T cells stimulated in vitro for 0 hrs or 24 hrs with or without IL-2 blockade. **(C)** Significant differentially expressed genes (DEG) (FDR cutoff <0.25) between Sh2b3^+/+^ and Sh2b3^-/-^ OT-I CD8+ T cells analyzed contrasts between each condition and genotype. Counts for DEGs were root-mean-squared scaled. DEGs were clustered by complete linkage hierarchical clustering and visualized by heat map. **(D)** Matrix visualization for genes of interest based upon bulk RNAseq transformed counts. (**E**) Pathway enrichment analysis using hallmark genesets filtered (FDR < 0.25, >5 DEG per pathway). The size per bubble corresponds to number of DEGs per pathway and coloring indicates the -log10 FDR values (*n* = 4 for 0 hr and 24 hr + anti-IL-2 conditions, *n* = 3 for 24 hr condition where *n* reflects individual mice).

Comparing the transcriptome at 24 hours of antigen stimulation, we identified DEG clusters of that normalized with anti-IL-2 treatment (**Fig. 4C-D**). Consistent with augmented surface expression of CD25 post-activation, we observed enhanced hallmark IL-2/STAT5-driven genes (*e.g. Cish, Eomes, Il2ra*) that normalized with blocking IL-2 (**Fig. 4D; Supplementary Fig. 5E)**. In addition, OT-I.*Sh2b3^-/-^* T cells exhibited enhanced terminal effector gene expression (*Ifng, Gzmb*, *Grzmc*) and augmented IL-12 receptor subunit expression (*Il12rb1, IL12rb2*). Blocking autocrine IL-2 normalized the STAT5 and MYC signature across genotypes, while the IFN-stimulated gene signature was elevated in SH2B3-deficient T cells (**Fig. 4E**). Based upon these findings, we concluded that SH2B3 negatively regulates autocrine IL-2 signaling in naïve CD8+T cells, leading to augmented STAT5 signaling and effector program following antigen stimulation.

### SH2B3 protects against diabetes in an induced T1D murine model

Based on the above observations, we hypothesized that enhanced effector function in SH2B3-deficient CD8+ T cells would exacerbate T1D in vivo. To test this idea, we utilized the RIP-mOVA mouse model, where membrane-bound ovalbumin (mOVA) is specifically expressed by pancreatic β-cells. In this model, adoptive transfer of OVA-specific OT-I cells (C57BL/6 background) has led to T1D development only with co-transferred, OVA-specific, CD4+ T cell help^38^. We isolated and transferred naïve CD8+ T cells from OT-I.*Sh2b3^+/+^* or OT-I.*Sh2b3^-/-^* into RIP-mOVA^+^ recipients in the absence of T cell help (**Fig. 5**). In contrast to recipients of OT-I.*Sh2b3^+/+^*T cells, all recipients of OT-I.*Sh2b3^-/-^* T cells developed diabetes starting at 7 days (**Fig. 5A-B**). Comparing the histologic features of pancreatic insulitis between genotypes, pancreata from recipients of the highest dose of *Sh2b3^+/+^* T cells demonstrated only low grade (1-2) insulitis with retention of insulin expression and only T cell infiltrates, when present, restricted to a peri-islet location (**Fig. 5C-D**). In contrast, pancreata from recipients of *Sh2b3^-/-^* T cells demonstrated high grade (3-4) insulitis with diffuse T and B cell infiltrates and mostly fibrotic insulin-negative islets by day 10 (**Fig. 5D**). Insulitis was not detected in recipients of either T cell genotype at day 3, but present at day 5 in hosts receiving OT-I.*Sh2b3^-/-^* T cells (**Supplementary Fig. 6A**). At day 3, SH2B3-deficient OT-I cells were already numerically expanded in the spleen of RIP-mOVA^+^ recipients compared to wildtype OT-I cells despite similar levels of proliferation (**Fig. 5F,G**). Therefore, cell intrinsic SH2B3 deficiency in self-reactive CD8+ T cells results in evasion of normal peripheral tolerance mechanisms.

**Figure 5.**
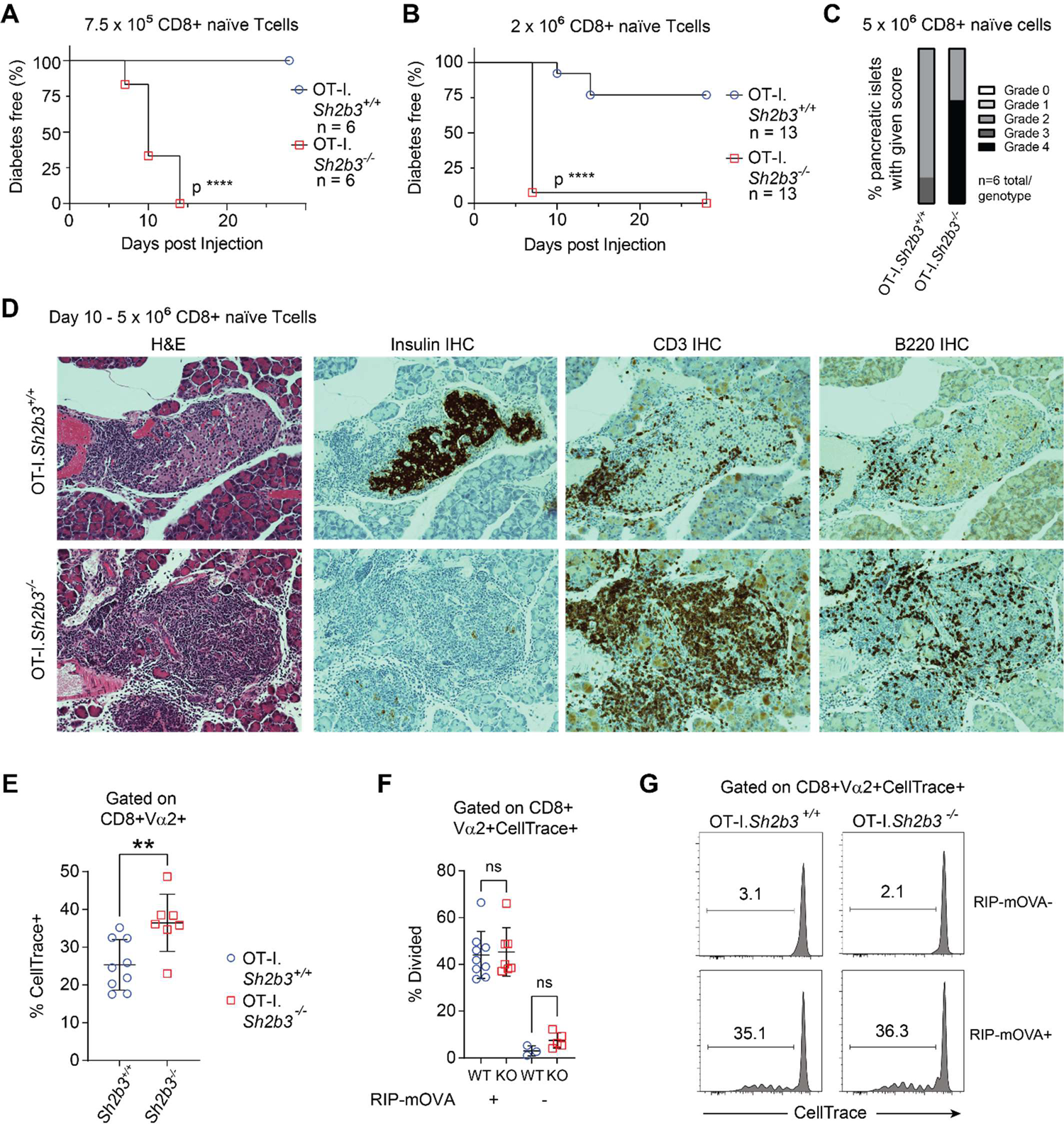
Adoptive transfer of SH2B3 -/- naïve OT-I CD8+ T cells initiates T1D development in RIP-mOVA recipient mice. (**A-G**) Purified naïve (CD44^lo^CD62L^hi^) from CD8+ OT-I.*Sh2b3^+/+^* or OT-I.*Sh2b3^-/-^* mice were adoptively transferred, at indicated cell dose, into RIP-mOVA^+^ or (**F-G**) transgene negative control recipient mice. (**A-B**) Diabetes incidence (blood glucose > 300mg/dL) in RIP-mOVA^+^ recipient mice was measured at days 5, 7, 10, and 28 days. (**C**) At days 5-10 post-transfer matched pancreata from recipients of each T cell genotype were fixed in formalin and processed for histological assessment (haematoxylin and eosin: H&E). Islets were graded for insulitis severity using a scale of 0-4: grade 0, normal tissue; grade 1 (scant) to grade 4 (severe). (**D**) Representative pancreas histology H&E and immunohistochemistry for insulin, CD3, and B220 are shown. (**E-G**) Recipient spleens were analyzed at Day 3 post-transfer for (**E**) prevalence of transferred cells or (**F**) percent cells divided with representative histograms showing proliferation based upon CellTrace dilution in (**G**). Significance was calculated using the logrank/Mantel-Cox (****P < 0.0001) and unpaired t test (**P < 0.01).

### SH2B3-deficient CD8+T cells exhibit a competitive survival advantage in RIP-mOVA recipient mice

Based on the increased proportion of OT-I.*Sh2b3^-/-^* T cells following adoptive transfer, we next compared the T cell responses in a competitive setting. We intravenously co-transferred naïve OT-I cells from each genotype into the same RIP-mOVA recipients (**Fig. 6A**). To distinguish the donor types, we utilized the tdTomato (Ai14) reporter and congenic (CD45.1 or CD45.2) alleles to delineate host and donor cells. Purified naïve OT-I cells of each genotype (**Supplementary Fig. 6B**) were mixed 1:1 prior to co-transfer into RIP-mOVA hosts (**Fig. 6A**). At both 3- and 5-days post-transfer, the relative proportion of OT-I.*Sh2b3^flox/flox^* T cells was increased in both the spleen and pancreas compared to OT-I.*Sh2b3^+/+^*cells transferred into the same host (**Fig. 6B**). Notably, SH2B3-deficient T cells accumulated despite comparable proliferation peaks and equivalent proliferation indexes for both genotypes (**Fig. 6C,D**). The competitive advantage of OT-I.*Sh2b3^flox/flox^*T cells in the absence of increased proliferation support the concept that loss of SH2B3 promotes increased CD8+ T cell survival. The accumulation of SH2B3-deficient cells required antigen as the CellTrace was undiluted in RIP-mOVA^-^ hosts (**Supplementary Fig. 6C**). Overall, following adoptive transfer into RIP-mOVA mice, OT-I.*Sh2b3^-/-^* CD8+ T cells exhibit enhanced survival and mediate CD4-independent, enhanced cytotoxicity following antigen stimulation leading to exacerbated disease onset and severity compared to control OT-I cells.

**Figure 6.**
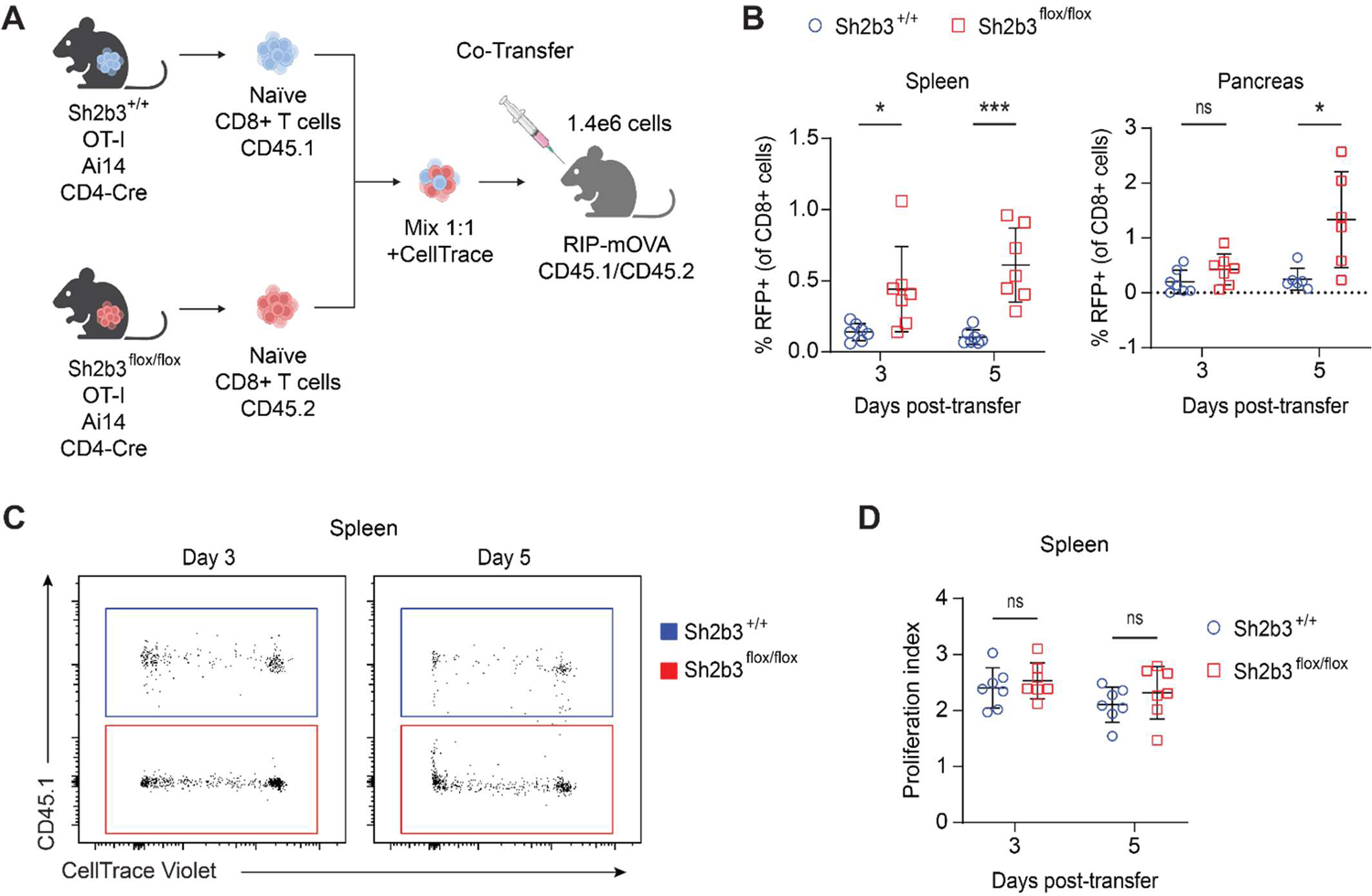
Competitive advantage for Sh2b3^-/-^ vs. Sh2b3^+/+^ OT-I CD8+ T cells in vivo in an identical RIP-mOVA+ host. (**A**) Experimental schematic. Naïve CD8+ T cells were purified from OT-I.*Sh2b3^+/+^*and OT-I.Sh2b3^flox/flox^ mice that were positive for CD4-Cre and Ai14 transgenes and disparate for the CD45 congenic marker (CD45.1 versus CD45.2). Naïve CD62L^+^CD44^-^CD8^+^ T cells were mixed 1:1 before being loaded with CellTrace Violet and transferred into RIP-mOVA+ hosts at 1.4e6 cells/host (7e5/genotype). (**B**) Graphs depict the proportion of donor OT-I.Sh2b3^+/+^ and OT-I.Sh2b3^-/-^ RFP^+^CD8^+^ T cells as a percentage of total CD8+ cells in the spleen and pancreas of RIP-mOVA^+^ hosts 3- and 5-days post-transfer. (**C**) Plots depict representative CellTrace Violet histograms of OT-I.Sh2b3^+/+^ and OT-I.Sh2b3^-/-^ CD8+ T cells in RIP-mOVA^+^ and RIP-mOVA^-^ hosts at 3 and 5 days post-transfer. (**D**) Graphs depict the proliferation index of donor OT-I.Sh2b3^+/+^ and OT-I.Sh2b3^-/-^ cells isolated from the spleen of RIP-mOVA^+^ hosts at 3 and 5 days post-transfer. Statistical significance was calculated using two-tailed t tests with corrects for multiple hypotheses in GraphPad Prism (A – B) (*n* = 6 for the 3 day transfer over two independent experiments, *n* = 3 for the 5 day transfer where *n* reflects the number of individual RIP-mOVA^+^ hosts) (C – E) (*n* = 7 over two independent experiments where *n* reflects the number of individual RIP-mOVA^+^ hosts). *P < 0.05, ***P < 0.001.

### Diabetes development is accelerated by SH2B3 deficiency in NOD mice

To test SH2B3 deficiency in a more physiologically relevant diabetes model, we created global SH2B3 deficient *NOD*/ShiLtJ (NOD) mice. NOD mice parallel spontaneous human T1D with numerous shared genetic alleles^39^. However, NOD mice lack an orthologous SH2B3 risk variant^40^. Therefore, we introduced the *Sh2b3^-/-^* loci from C57BL/6 mice onto the NOD genetic background by backcrossing prior to intercrossing for littermate control cohorts utilized in diabetes incidence studies. NOD mice spontaneously develop pancreatic insulitis starting at 5 weeks of age followed by the development of diabetes typically by 10-12 weeks^41^. In our NOD colony, ∼80-85% of female mice develop diabetes, while ∼50% of male mice develop diabetes over the course of 35 weeks. Here, we found that introduction of the *Sh2b3* knockout allele on the NOD background resulted in accelerated diabetes in both female and male mice (**Fig. 7**). Across sexes, the NOD.*Sh2b3^-/-^* mice developed diabetes at an accelerated rate relative to their NOD.*Sh2b3^+/+^* counterparts. In male NOD mice, diabetes development was more pronounced in SH2B3 deficient mice, with heterozygous littermates exhibiting an intermediate phenotype. Collectively, our results confirm a protective, dose-dependent, effect of SH2B3 against development of diabetes across alternative murine models of T1D.

**Figure 7.**
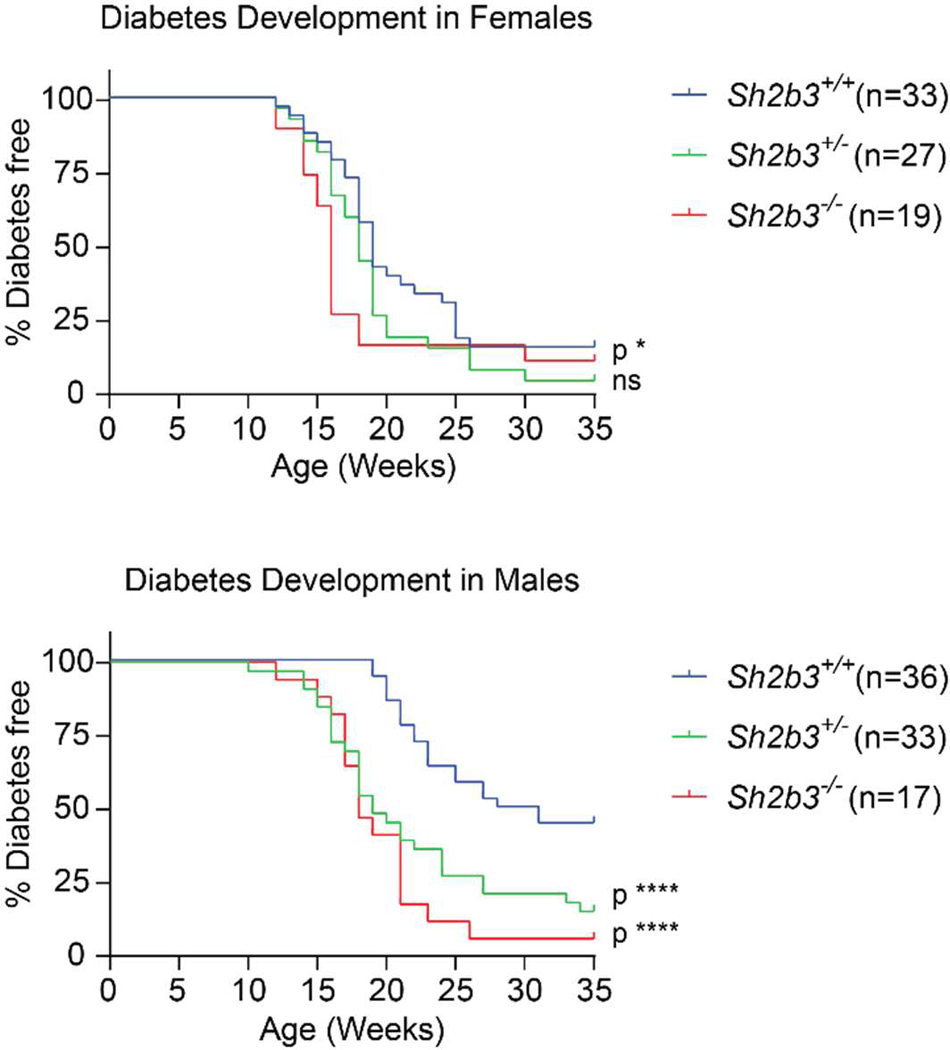
Loss of SH2B3 function promotes diabetes development in a spontaneous T1D murine model. Sh2b3^-/-^ mice were backcrossed with NOD/ShiLtJ mice for 15 generations prior to intercrossing to generate NOD.Sh2b3^-/-^ mice. Littermate control cohorts were monitored for diabetes development based on blood glucose levels. Graphs depict the prevalence of diabetes NOD.*Sh2b3^+/+^*, NOD.*Sh2b3^+/-^*, and NOD.*Sh2b3^-/-^* mice over the course of 35 weeks, separated based on sex. Significance was calculated using the Gehan-Breslow-Wilcoxon test in GraphPad Prism. *P < 0.05, ****P < 0.0001.

## DISCUSSION

Here, we demonstrate that reduced function of the adaptor protein SH2B3 contributes to a loss of peripheral T cell tolerance and autoimmune diabetes in both humans and mice. First, we identified a single *SH2B3* haplotype significantly associated with increased risk for T1D, and this haplotype carries the rs3184504*T missense variant encoding a hypomorphic SH2B3^262W^ protein^9^. In TEDDY studies in children with high-risk HLA alleles^2^, the same autoimmune SH2B3^RISK^ haplotype was identified as one of the few genetic risk loci predictive of anti-islet autoantibodies^42^. We performed functional studies in murine models and found a critical role for SH2B3 in maintaining peripheral T cell tolerance to pancreatic β-cell antigens. Herein, we used the well-established RIP-mOVA mouse model and found OVA-specific OT-I.*Sh2b3^-/-^*T cells caused early islet destruction and diabetes without requiring CD4+ T cell help. In contrast, control self-reactive OT-I cells did not induce diabetes. Transferred SH2B3-deficient cells were more abundant than their wildtype counterparts even in the same recipient environment, yet they showed comparable proliferation, suggesting a cell-intrinsic survival advantage. Finally, we created a NOD*.Sh2b3^-/-^* spontaneous diabetes model demonstrating that global deficiency of SH2B3 in NOD mice led to increased incidence and accelerated diabetes across sexes compared to wild-type NOD mice. Taken together, these data suggest that SH2B3 has an underappreciated T cell intrinsic role in peripheral tolerance and that both global and T cell specific reduction of SH2B3 functionality contributes to T1D.

SH2B3 is an inducible adaptor protein that binds to phosphorylated JAK2 and JAK3 regulating the intensity and duration of cytokine responses^10,17^. In this study, we identified a T cell-intrinsic role for SH2B3 in regulating γ_c_ and IFN-γ cytokine signaling, including the IL-2R feedback loop early post-TCR stimulation (**Fig. 8**). We observed *SH2B3* upregulation in T lymphocytes within hours of antigen stimulation both in human and mouse cells. Additionally, in the first days following antigen encounter, SH2B3-deficient cells upregulated CD25 more than control T cells and showed strong transcriptional enrichment in IL-2/STAT5, MYC, and mTORC1 gene sets, the majority of which normalized when IL-2 was blocked in culture. Importantly, IL-2 production was not affected by SH2B3 deficiency. Unlike many negative regulators in T cells, the loss of SH2B3 had no impact on TCR signaling strength, as shown by lack of changes in CD69 expression, immediate early gene transcripts, or T cell proliferation. Consistent with previous reports, we found SH2B3-deficiency had little impact on thymocyte development or peripheral naïve T cell homeostasis. Strikingly, differential CD25 upregulation in *Sh2b3^-/-^*vs *Sh2b3^+/+^* T cells in response to TCR engagement was most evident in the setting of low-affinity peptides, suggesting the impact of reduced SH2B3 function is most critical in the context of limiting antigen or lower-affinity TCRs, such as those expressed by self-reactive T cells.

**Figure 8.**
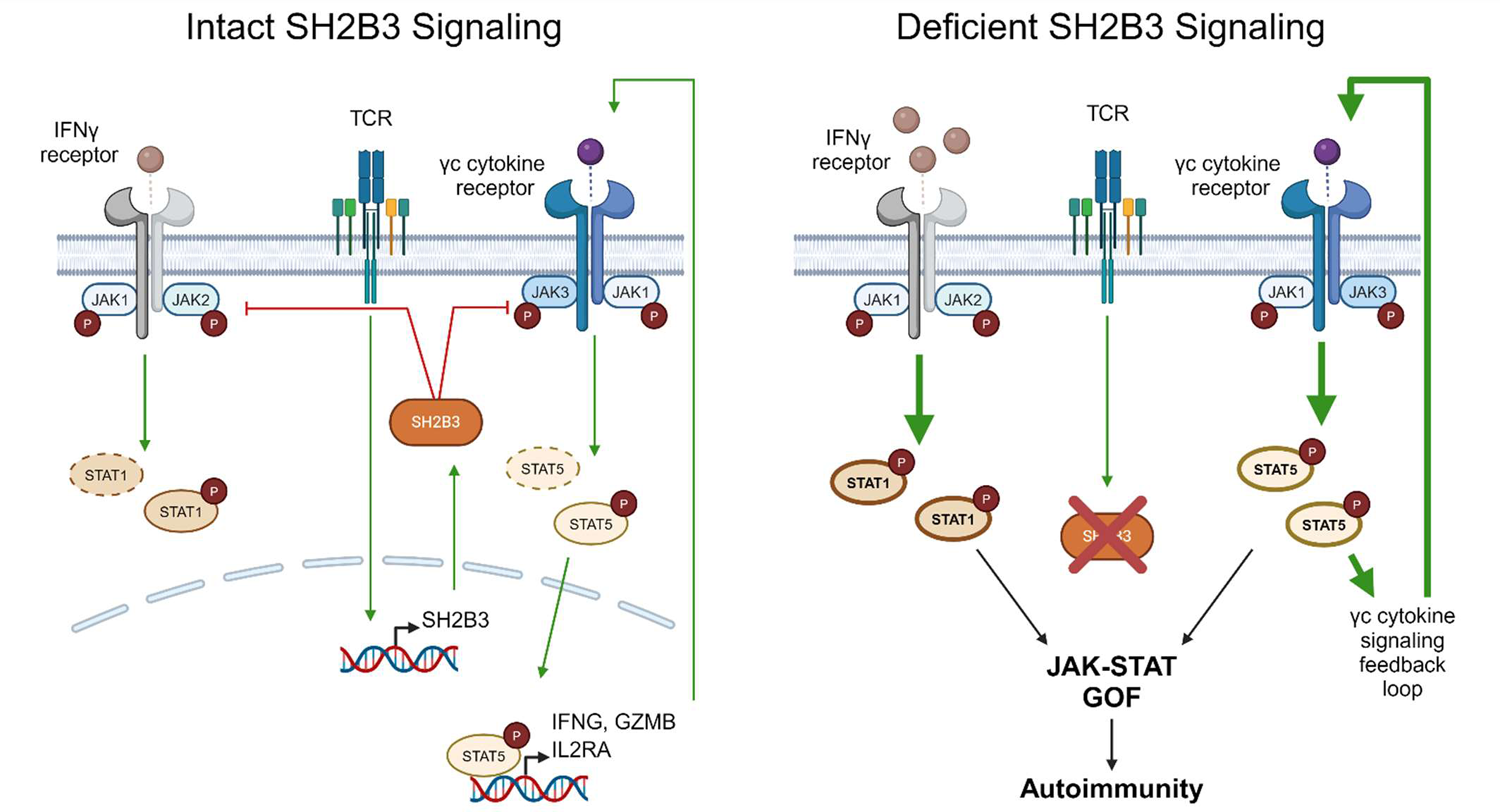
Schematic of SH2B3 regulation of cytokine and TCR signaling in T cells. *SH2B3* expression is upregulated following TCR stimulation. SH2B3 directly negatively regulates JAK1/JAK3 phosphorylation of STAT5 downstream of γc cytokine signaling as well as JAK1/JAK2 phosphorylation of STAT1 downstream of IFN-γ signaling. This negative regulation helps maintain peripheral tolerance of autoreactive T cells. With disrupted SH2B3 function, STAT1 and STAT5 phosphorylation is increased. This results in a positive feedback loop in γc cytokine signaling pathways, promoting a JAK-STAT gain of a function phenotype. Hyperactive JAK-STAT signaling increases the ability of autoreactive T cells to evade peripheral tolerance, contributing to the development of T1D and other autoimmune conditions.

Consistent with strong early IL-2 signaling, OT-I.*Sh2b3*^-/-^ cells also showed increased expression of effector molecules *(Ifng, Gzmb, Prf1*) compared to wildtype cells^43^. Lastly, we found enhanced survival of *Sh2b3^-/-^* CD8+ T cells compared to *Sh2b3^+/+^*T cells following antigen exposure. Strong initial IL-2 signaling observed in SH2B3 deficient cells also likely synergizes with additional cytokine signaling. Previous reports showed augmented IL-15 signaling in memory *Sh2b3^-/-^* CD8^+^CD44^+^ T cells^35^. We and others have also shown enhanced IL-7 signaling in *Sh2b3^-/-^* B cell progenitors^30,44^. Additionally, SH2B3 regulates JAK-STAT signaling downstream of the Th1-related cytokine IL-12^45^. Herein, we found SH2B3 deficient T cells showed hypersensitivity across subsets to IL-2, IL-7, IL-15. and IFN-γ, depending upon the cell- and activation-state that alter cytokine receptor expression. Collectively, our data demonstrate that SH2B3 functions post-activation in CD8+ T cells to limit sensitivity to several JAK2/3 regulated pathways to further shape differentiation and survival.

Peripheral CD8+ T cell tolerance is a major checkpoint preventing T1D. In our work, we found SH2B3 deficient OT-I cells were resistant to peripheral tolerance mechanisms in RIP-mOVA hosts showing enhanced survival and robust effector function. Previous work using the RIP-OVA model identified a CD8+ “tolerant” phenotype characterized by low effector molecule and cytokine receptors (IL2RA, IL7R) expression that can be reversed by a strong TCR and inflammatory signal (immunization or infection with cognate antigen)^46,47^. Breach in tolerance correlated with a strong early IL-2-STAT5/MYC expression, enhanced effector phenotype and altered metabolism. Similarly, another group showed that enhanced early IL-2 signaling in OT-I cells from Treg-depleted RIP-OVA hosts promoted diabetes and a distinct CD8^+^IL-7R^+^ effector cell with superior cell-killing abilities^36^. Herein, we found the synergistic effect of hypersensitivity to many γ_c_ and IFN-γ cytokine pathways in SH2B3 deficiency potently promotes pathogenic CD8+ T cells. To model the impact of global SH2B3 deficiency on spontaneous diabetes, we created the NOD.*Sh2b3^-/-^*mouse and found high incidence and accelerated diabetes across sexes compared to control NOD animals. Many cell types contribute to insulitis and diabetes in the NOD model, including CD4+ effector, Tregs, islet cells and myeloid cells. In our previous work, we found reduced SH2B3 augmented myelopoiesis aiding in the protection from sepsis^20^, but these changes in cell infiltration limited tissue pathology. In NOD mice, CD4+ T cells initiate pancreatic infiltration and provide help to cytotoxic T cells, while CD8+ T cells are primarily responsible for β-cell destruction^48^. Herein, we demonstrate that both SH2B3 deficient CD4+ and CD8+ T cells demonstrate enhanced cytokine responsiveness including increased IL-2R signaling following TCR stimulation. Thus, it is tempting to speculate that loss of SH2B3 in both CD4+ and CD8+ effector T cells contributes to the development of diabetes in our NOD model. Additionally, the presence of autoreactive T cells is not sufficient for diabetes as boosting Treg function and numbers can prevent islet destruction^49^. As expected with enhanced IL-2R signaling, we found an expanded Treg population in global *Sh2b3^-/-^*mice. Similarly, Zhang et al found in mice with a reduced function Sh2b3 knockin allele there was an increased Treg population yet similar in vitro suppressive function compared to wildtype cells^16^. Additionally, certain autoimmune risk genotypes can render effector T cells resistant to suppression^38,50^. For example, PTPN2-deficient T cells from NOD mice also were resistant to Treg suppression despite the mice having increased Treg frequency^38^. Despite the increased proportion of Treg in *Sh2b3^-/-^* mice, global NOD.*Sh2b3^-/-^* mice exhibited an increased incidence of diabetes, thus testing the ability of effector T cells to be suppressed will be important future experiments. Overall, we found loss of SH2B3 exacerbated diabetes in both the RIP-mOVA and the spontaneous NOD.*Sh2b3^-/-^* mouse models.

In summary, our data show that reduced SH2B3 function leads to loss of T cell tolerance and suggest an important T-cell intrinsic regulatory role for SH2B3 in T1D. Our findings support the hypothesis that the SH2B3^262W^ risk protein is a causal variant for T1D and likely multiple other autoimmune diseases. Future work using the models reported here will allow for further mechanistic studies on how SH2B3 contributes to human autoimmunity.

## ACKNOWLEDGEMENTS

We thank Jit Khim and Anna Zielinska-Kwiatkowska for their maintenance and care of animals.

## FUNDING

This study was funded by the National Institute of Diabetes and Digestive and Kidney Diseases (DP3DK111802, K08DK114568, and R03DK134746).

## AUTHOR CONTRIBUTIONS

T.K.W., A.B.I.R., T.D., and E.J.A. designed and performed experiments, analyzed data, and wrote and/or edited the manuscript; J.A.M., M.A.M., and H.D.L. developed required models, strains, or reagents and/or performed experiments; Y.G., P.C., and J.H.B. designed and interpreted human subject studies; D.A.G., K.A.D., and M.C.A. performed bioinformatic analysis; D.J.R. and E.J.A. conceived of and supervised the study, interpreted data, and edited the manuscript.

## COMPETING FINANCIAL INTERESTS

The authors declare no competing financial interests.

## Notes

### Competing Interest Statement

The authors have declared no competing interest.

