## Supplementary material for "Reduced function of the adaptor SH2B3 promotes T1D via altered gc cytokine-regulated, T cell intrinsic immune tolerance": Online Supplemental Material

**1. Supplementary Table I.** Single-marker family-based association analysis of variants in *SH2B3* and *ATXN2*

**2. Supplementary Table I.** Single-marker family-based association analysis of variants in *SH2B3* and *ATXN2*

**3. Supplementary Figure 1.** *Sh2b3*$^{-/-}$ T cells lack SH2B3 protein expression and exhibit increased STAT5 phosphorylation following stimulation

**4. Supplementary Figure 2.** *Sh2b3* deficiency doesn't significantly impact T cell development and minimally impacts circulating T cell populations

**5. Supplementary Figure 3.** Sh2b3 deficient T cells exhibit increased cytokine sensitivity with minimal differences at baseline

**6. Supplementary Figure 4.** Sh2b3 deficient cells exhibit increased CD25 expression following TCR stimulation with equal IL-2 concentration

**7. Supplementary Figure 5.** SH2B3 function minimally impacts transcriptional regulation in the early stages of TCR activation

**8. Supplementary Figure 6.** *Sh2b3*$^{-/-}$ naïve OT-I CD8$^{+}$ T cells cause accelerated islet destruction and fibrosis in RIP-mOVA recipients

**9. Supplementary Methods**

**10. Supplementary References**

**Supplemental Table I.** Single-marker family-based association analysis of variants in *SH2B3* and *ATXN2*

| dbSNP ID (137) | Location | cDNA change | Amino Acid change | MAF | Family (n) | *Z* score | *P* value |
|---|---|---|---|---|---|---|---|
| **rs3184504** | **Exon 3 (SH2B3)** | **c.784T>C[a]** | **p.Trp262Arg** | **0.475** | **1423** | **-6.639** | **$3.23 \times 10^{-11}$** |
| **rs653178** | **Intron 1 (ATXN2)** | **c.732-14033G>A[b]** | **None** | **0.474** | **1422** | **-6.823** | **$9.09 \times 10^{-12}$** |

Single-marker analysis in T1D-affected sibling pairs and trio families of European ancestry ascertained by the Type 1 Diabetes Genetics Consortium. The location, cDNA change and amino acid change for rs3184504 (GRCh38 NC_000012.12:g.111446804T>C) are based on the RefSeq sequences NM_005475.3 and NP_005466.1. The location and cDNA change for rs653178 (GRCh38 NC_000012.12:g.111569952C>T) are based on the RefSeq sequence NM_002973.3. Z scores and P values are calculated for the minor alleles. Positive and negative Z scores indicate risk and protection, respectively.

Abbreviations: MAF, minor allele frequency; n, number of informative families.

[a]For rs3184504, C is the minor allele in this cohort, whereas the T is the minor allele in the 1000Genomes European population.
[b]For rs653178, A is the minor allele in this cohort, whereas the G is the minor allele in the 1000Genomes European population.

**Supplemental Table II.** Family-based haplotype association analysis of *SH2B3/ATXN2*

| **Haplotype rs3184504/rs653178** | **Frequency** | **Family (n)** | ***Z* score** | ***P* value** |
|---|---|---|---|---|
| **H1 T (p.262W)/G** | **0.532** | **1358** | **6.630** | **$3.42 \times 10^{-11}$** |
| **H2 C (p.R262)/A** | **0.465** | **1361** | **-6.677** | **$2.48 \times 10^{-11}$** |
| **H3 C (p. R262)/G** | **0.002** | **18** | **1.254** | **0.209667** |
| **H4 T (p.262W)/A** | **0.001** | **5** | **-1.667** | **0.095581** |

Haplotype analysis of *SH2B3* in T1D-affected sibling pairs and trio families of European ancestry (quantification, column 3) ascertained by the Type 1 Diabetes Genetics Consortium. Positive and negative *Z* scores indicate risk and protection, respectively.

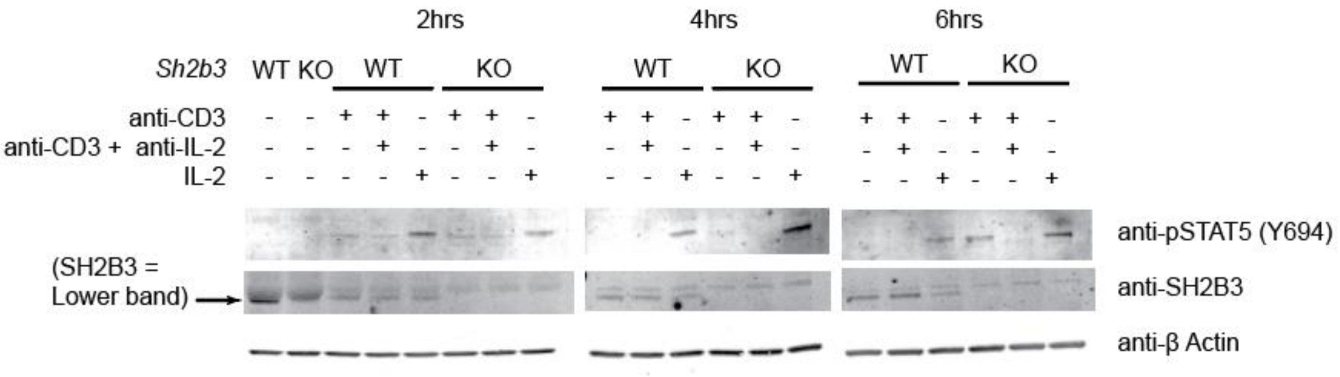


**Supplementary Figure 1. *Sh2b3*$^{-/-}$ T cells lack SH2B3 protein expression and exhibit increased STAT5 phosphorylation following stimulation.** SH2B3 expression and STAT5 phosphorylation in T cells following TCR and cytokine stimulation with β actin expression as a control. Naïve OT-I+ Sh2b3$^{+/+}$ and Sh2b3$^{-/-}$ were stimulated with murine IL-2 alone (100 ng/mL) or stimulated with anti-CD3 (10 ug/mL) with and without anti-IL-2 blocking antibody (10 ug/mL) for 2, 4, and 6 hrs.

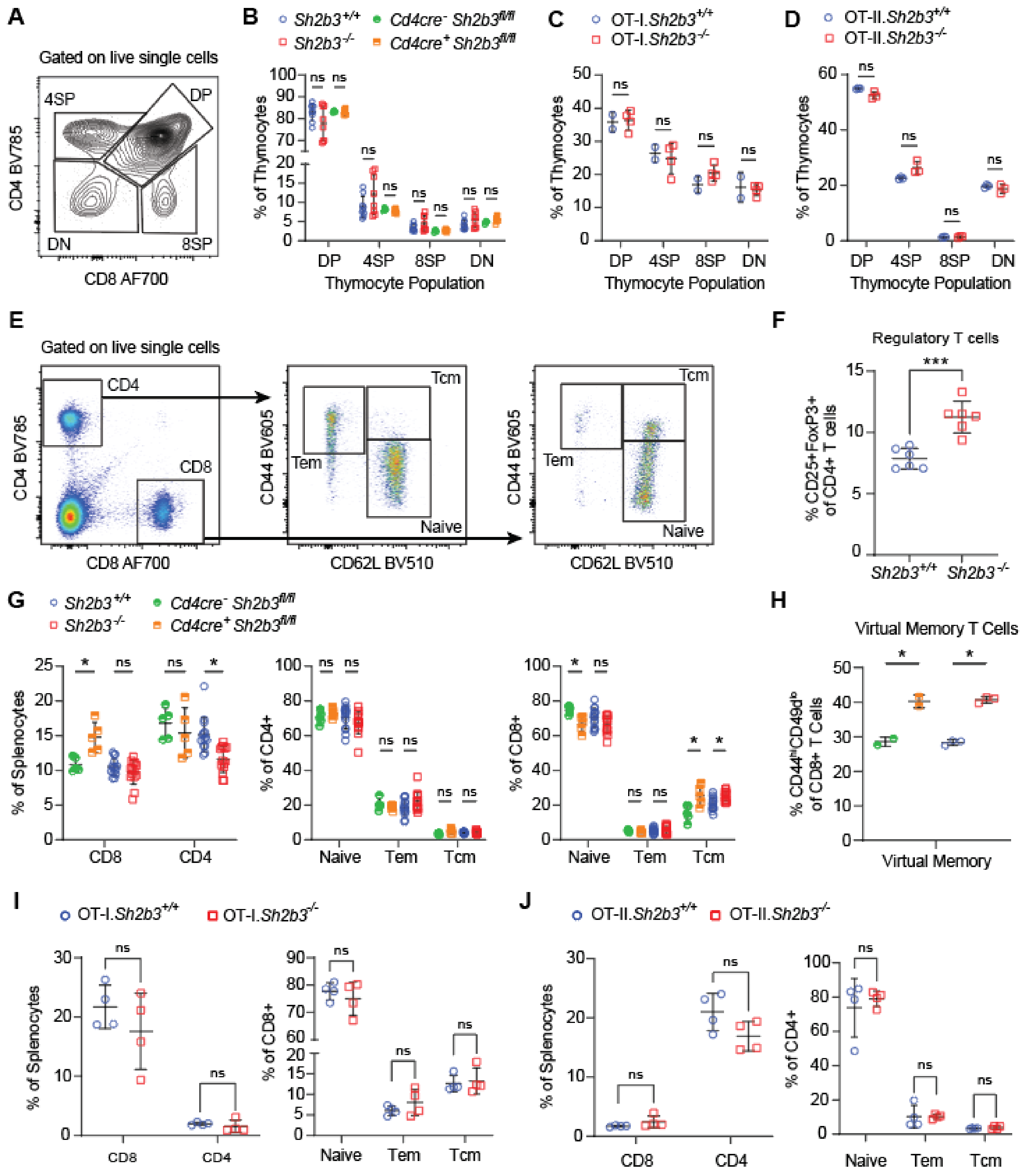


**Supplementary Figure 2. Sh2b3 deficiency doesn’t significantly impact T cell development and minimally impacts circulating T cell populations.** (**A**) Representative plot depicts the flow cytometry gating scheme phenotyping thymocytes. (**B** - **D**) Graphs depict the proportion of thymocytes that are DP, 4SP, 8SP, and DN from (B) polyclonal CD4-Cre$^{-}$Sh2b3$^{flox/flox}$, CD4-Cre$^{+}$Sh2b3$^{flox/flox}$, Sh2b3$^{+/+}$, and Sh2b3$^{-/-}$ mice, (C) OT-I.Sh2b3$^{+/+}$ and OT-I.Sh2b3$^{-/-}$ mice, and (D) OT-II.Sh2b3$^{+/+}$ and OT-II.Sh2b3$^{-/-}$ mice. (**E**) Representative plots depict the flow cytometry gating scheme of phenotyping splenocytes. (**F**) Graph depicts the proportion of CD4 single positive T cells in the spleen that are CD25+FoxP3+ in polyclonal Sh2b3$^{+/+}$ and Sh2b3$^{-/-}$ mice. (**G**) Graphs depict the proportion of splenocytes that are CD8 and CD4 single positive and, of the single positive populations, proportions that are naïve, Tem, and Tcm in polyclonal CD4-Cre$^{-}$Sh2b3$^{flox/flox}$, CD4-Cre$^{+}$Sh2b3$^{flox/flox}$, Sh2b3$^{+/+}$, and Sh2b3$^{-/-}$ mice. (**H**) Graph depicts the proportion of CD8 single positive T cells in the spleen that are antigen inexperienced virtual memory cells in polyclonal CD4-Cre$^{-}$Sh2b3$^{flox/flox}$, CD4-Cre$^{+}$Sh2b3$^{flox/flox}$, Sh2b3$^{+/+}$, and Sh2b3$^{-/-}$ mice. These virtual memory cells are defined as CD8$^{+}$ cells that are CD44$^{hi}$CD49d$^{lo}$. (**I**) Graphs depict the proportion of splenocytes that are CD8 and CD4 single positive and, of the CD8+ population, proportions that are naïve, Tem, and Tcm in OT-I.Sh2b3$^{+/+}$ and OT-I.Sh2b3$^{-/-}$ mice. (**J**) Graphs depict the proportion of splenocytes that are CD8 and CD4 single positive and, of the CD4+ population, proportions that are naïve, Tem, and Tcm in OT-II.Sh2b3$^{+/+}$ and OT-II.Sh2b3$^{-/-}$ mice. (B – D) (F – I) Significance was calculated using two-tailed unpaired t-tests in GraphPad Prism with corrections for multiple hypotheses. (B) $n$ = 3 for Sh2b3$^{flox/flox}$ mice, $n$ = 9 for Sh2b3$^{+/+}$ mice, and $n$ = 8 for Sh2b3$^{-/-}$ mice, (C) $n$ = 2 for OT-I.Sh2b3$^{+/+}$ mice, $n$ = 4 for OT-I.Sh2b3$^{-/-}$, (D) $n$ = 3, (F) $n$ = 6, (G) $n$ = 3 for Sh2b3$^{flox/flox}$ mice, $n$ = 13 for Sh2b3$^{+/+}$ mice and Sh2b3$^{-/-}$ mice, (H) $n$ = 2 for Sh2b3$^{flox/flox}$ mice, $n$ = 3 for Sh2b3$^{+/+}$ mice and Sh2b3$^{-/-}$ mice , and (I – J) $n$ = 4, where $n$ reflects individual mice. *P < 0.05, ***P < 0.001.

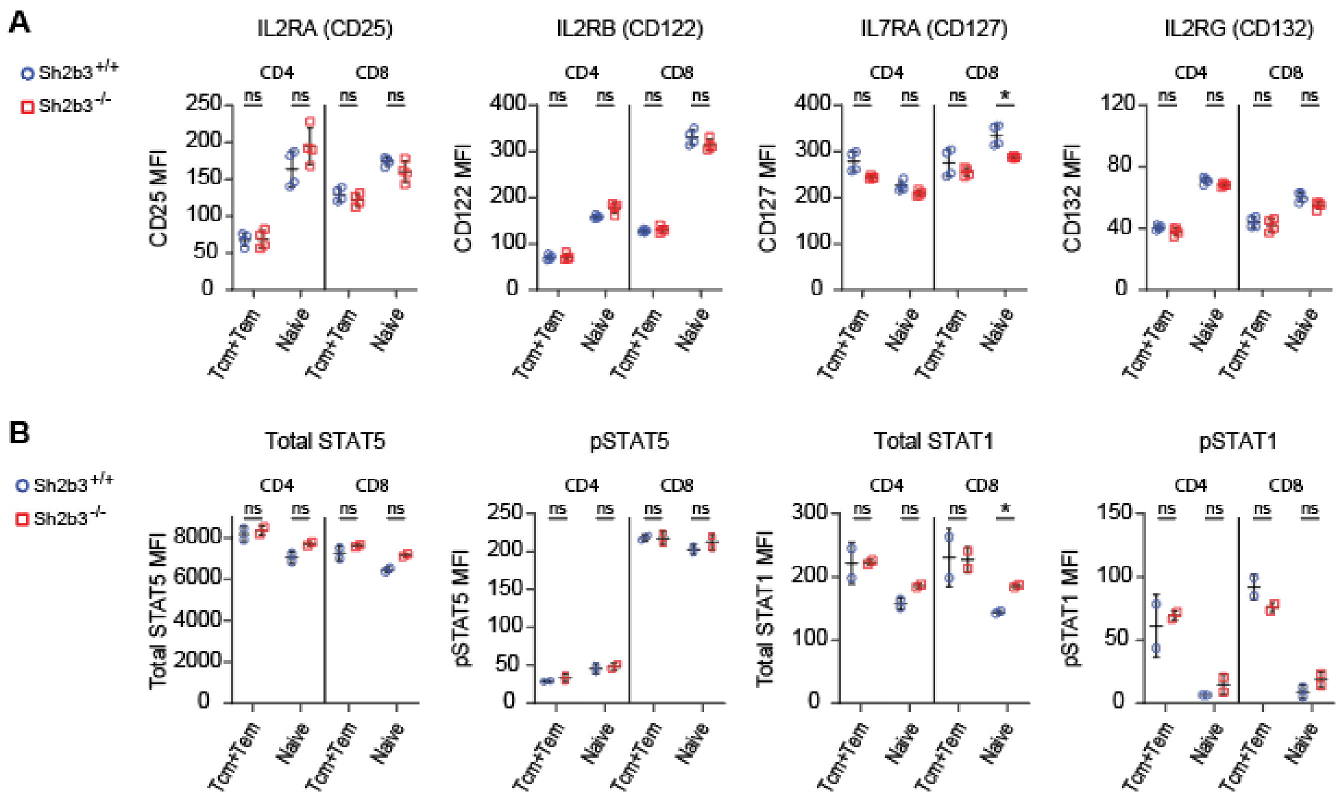


**Supplementary Figure 3. Sh2b3 deficient T cells exhibit increased cytokine sensitivity with minimal differences at baseline**. (**A**) Graphs depict the baseline expression of common gamma chain cytokine receptor subunits in naïve and memory CD4+ and CD8+ Sh2b3$^{+/+}$ and Sh2b3$^{-/-}$ cells. (**B**) Graphs depict the baseline expression of total and phosphorylated STAT1 and STAT5 in naïve and memory CD4+ and CD8+ Sh2b3$^{+/+}$ and Sh2b3$^{-/-}$ cells. Significance was calculated using two-tailed t-tests with corrections for multiple hypotheses in GraphPad Prism. (A) $n$ = 4, where $n$ reflects technical duplicates, (B) $n$ = 2, where $n$ reflects individual mice. *P < 0.05.

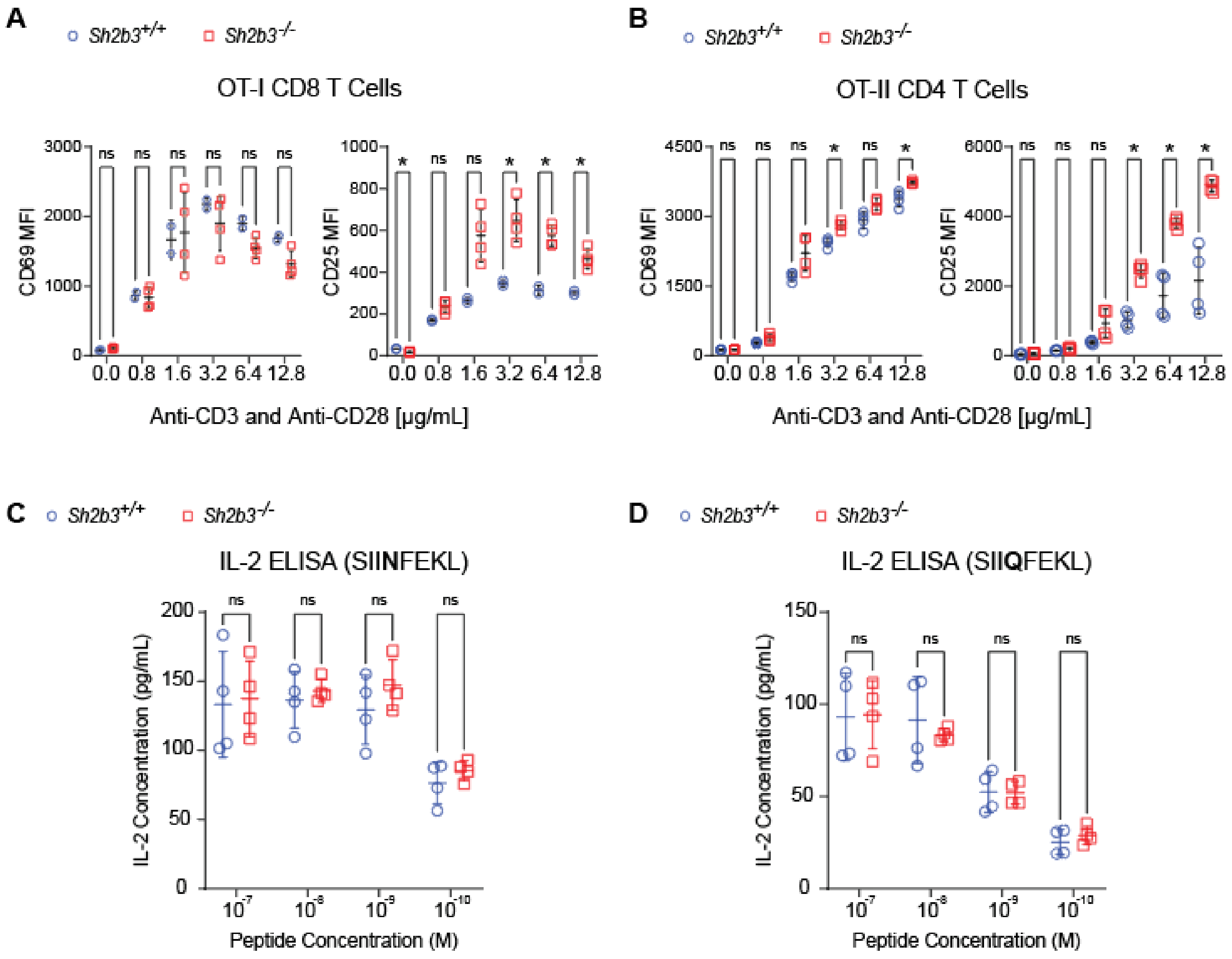


**Supplementary Figure 4. Sh2b3 deficient cells exhibit increased CD25 expression following TCR stimulation with equal IL-2 concentration.** (**A** - **B**) Graphs depict the MFI of CD25 and CD69 in (A) OT-I.Sh2b3$^{+/+}$ and OT-I.Sh2b3$^{-/-}$ naïve CD8+ T cells and (B) in OT-II.Sh2b3$^{+/+}$ and OT-II.Sh2b3$^{-/-}$ naïve CD4+ T cells after the indicated cells were plated at (A) 1e5/well and (B) 5e4/well in flat-bottomed 96-well plates with platebound anti-CD3 and anti-CD28 stimulatory antibodies at varying concentrations for 16 hrs of stimulation at 37C. (**C** - **D**) Graphs depict the concentration of IL-2 in media of wells holding CD8+ T cells stimulated with APCs loaded with SIINFEKL or SIIQFEKL. Naïve CD8+ T cells were plated at 7.5e4/well in U-bottomed 96-well plates with irradiated APCs (1e5/well) loaded with SIINFEKL altered peptide ligands (APLs) at $10^{-10}$ – $10^{-7}$ M for 72 hrs. (A – D) Statistical significance was calculated using two-tailed unpaired t-tests in GraphPad Prism with corrections for multiple hypotheses (*n* = 4, where *n* reflects technical duplicate stimulation wells) $^{*}P < 0.05$, $^{**}P < 0.01$.

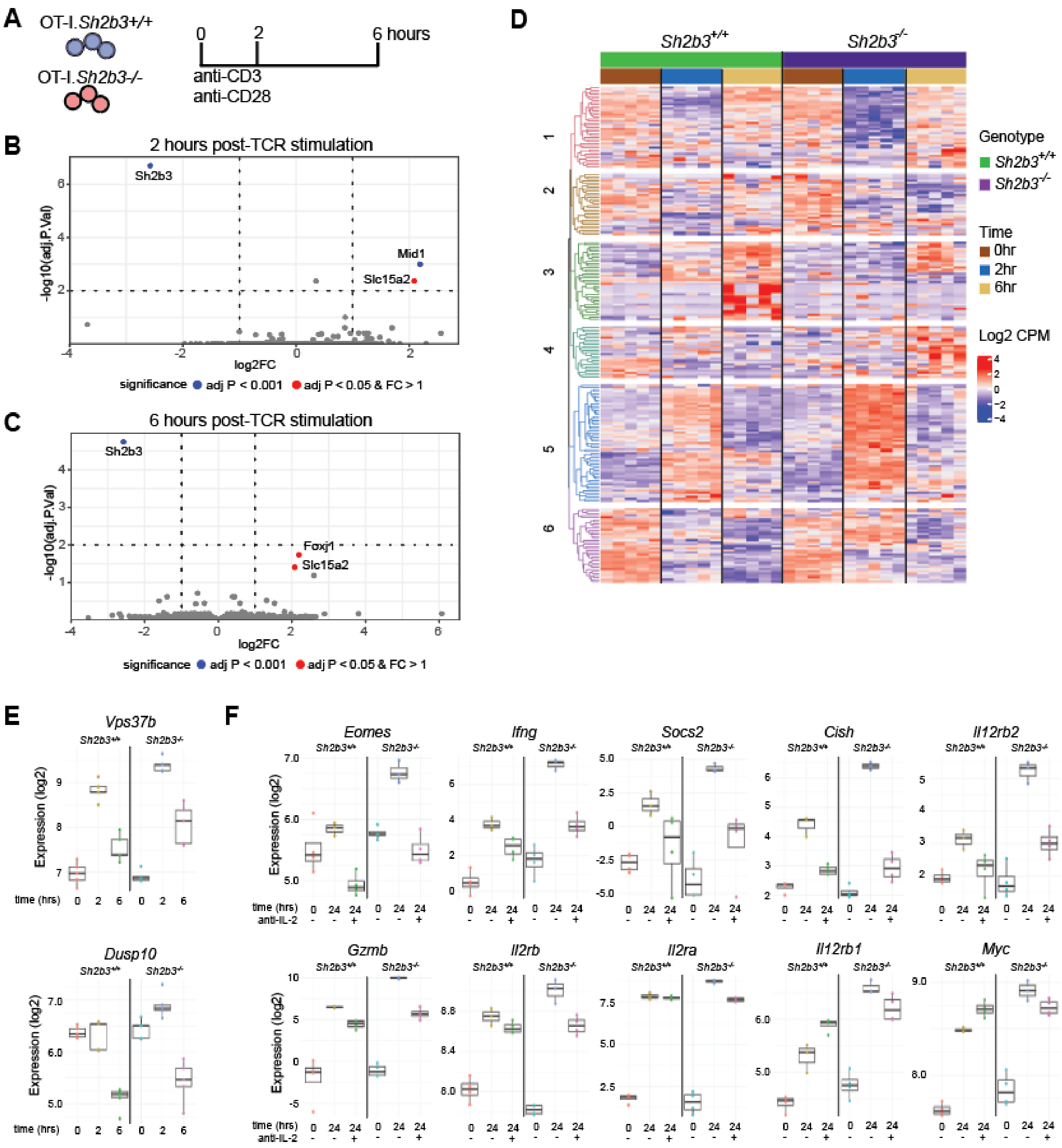


**Supplementary Figure 5. SH2B3 function minimally impacts transcriptional regulation in the early stages of TCR activation.** (**A**) Naïve CD8+ T cells were isolated from OT-I.Sh2b3$^{+/+}$ and OT-I.Sh2b3$^{-/-}$ mice and plated with 10 ug/mL anti-CD3 and 5 ug/mL anti-CD28 stimulatory antibodies for 0, 2, and 6 hours with and without 10ug/mL anti-IL-2 blocking antibody before RNA was isolated and sequenced. (**B** - **C**) Volcano plot depicts differentially expressed genes at (A) 2 hrs and (B) 6 hrs of TCR stimulation between OT-I.Sh2b3$^{+/+}$ and OT-I.Sh2b3$^{-/-}$ CD8+ T cells. Naïve CD8+ T cells were isolated from OT-I.Sh2b3$^{+/+}$ and OT-I.Sh2b3$^{-/-}$ mice and plated with 10 ug/mL anti-CD3 and 5 ug/mL anti-CD28 stimulatory antibodies. (**D**) Bulk RNAseq data on expression in naïve CD8+ T cells stimulated *in vitro* for 0 hrs, 2hrs, and 6 hrs. (**E**) Graphs depict expression of select genes of interest in OT-I.Sh2b3$^{+/+}$ and OT-I.Sh2b3$^{-/-}$ CD8+ T cells after 24 hours of stimulation with and without anti-IL-2 blocking antibody.

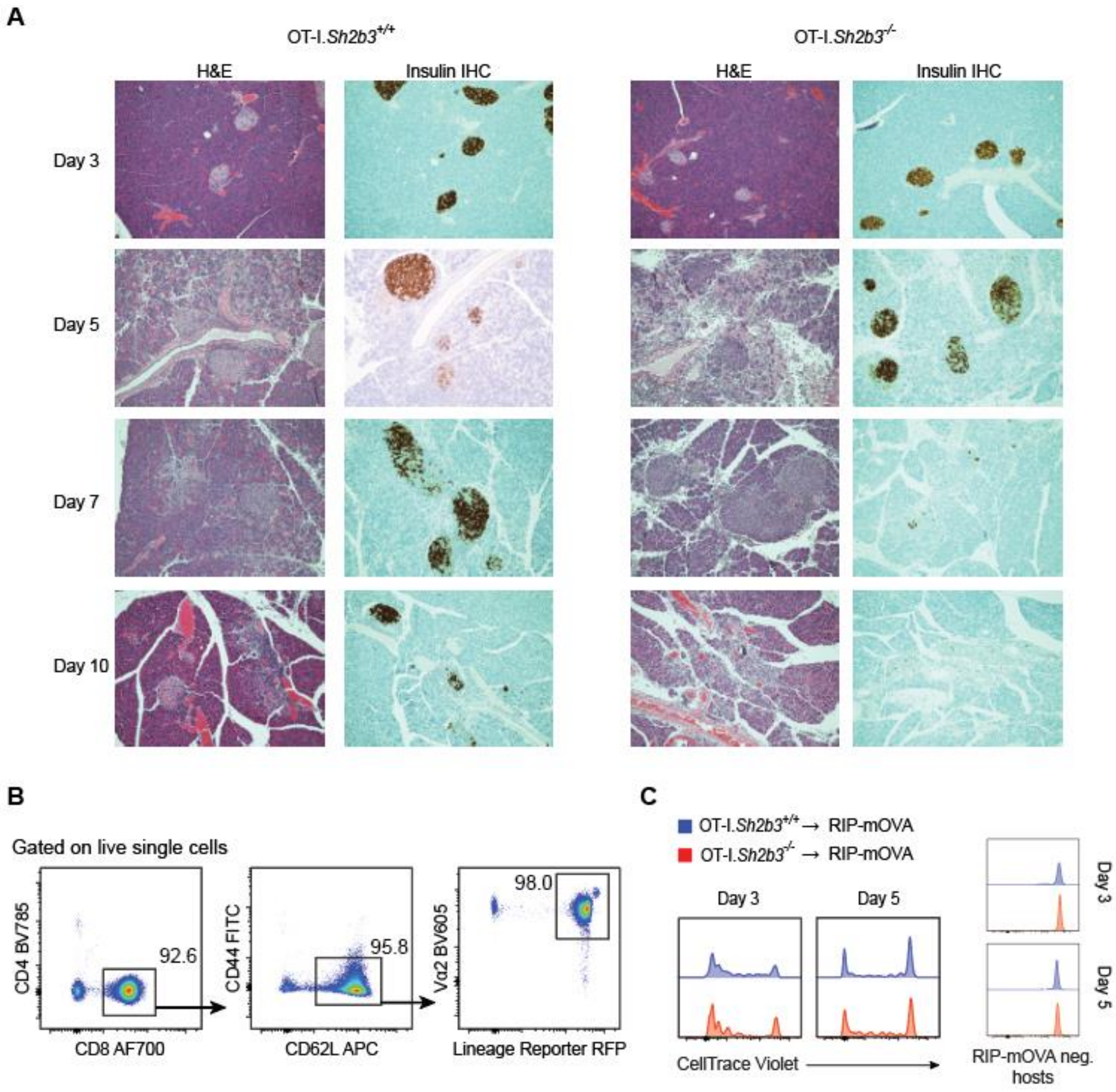


**Supplementary Figure 6. *Sh2b3*$^{-/-}$ naïve OT-I CD8$^{+}$ T cells cause accelerated islet destruction and fibrosis in RIP-mOVA recipients.** (**A**) Insulitis progression in RIP-mOVA mice at days 3, 5, 7, and 10 following transfer of 5e6 OT-I.*Sh2b3*$^{+/+}$ or OT-I.*Sh2b3*$^{-/-}$ naïve CD8$^{+}$ T cells. H&E and immunohistochemistry staining for insulin was performed. Patchy insulitis was shown in sections from mice that received wildtype cells and widespread early islet destruction in sections from mice receiving SH2B3-deficient T cells. Representative histology shown of n=3/genotype. (**B**) Representative plots depict the flow cytometry gating scheme to quantify the purity of CD8$^{+}$ CD62L$^{hi}$CD44$^{lo}$ RFP$^{+}$Vα2$^{+}$ cells being transferred into RIP-mOVA hosts. (**C**) Plots depict representative CellTrace Violet histograms of OT-I.Sh2b3$^{+/+}$ and OT-I.Sh2b3$^{-/-}$ CD8$^{+}$ T cells in RIP-mOVA$^{+}$ and RIP-mOVA$^{-}$ hosts at 3- and 5-days post-transfer of 1.5e6 naïve CD8$^{+}$ T cells.

### SUPPLEMENTARY MATERIALS & METHODS

| REAGENT | Manufacturer | Product # |
|---|---|---|
| **Antibodies** | | |
| BV785 anti-mouse CD4 (GK1.5) | BioLegend | #100453 |
| AF700 anti-mouse CD8a (53-6.7) | BioLegend | #100730 |
| BV605 anti-mouse/human CD44 (IM7) | BioLegend | #103047 |
| PerCP/Cyanine5.5 anti-mouse CD4 (GK1.5) | BioLegend | #100434 |
| PE/Cyanine7 anti-mouse CD69 (H1.2F3) | BioLegend | #104512 |
| Pacific Blue anti-mouse CD25 (PC61) | BioLegend | #102022 |
| FITC anti-mouse CD122 (TM-β1) | BioLegend | #123208 |
| PE anti-mouse FOXP3 (150D) | BioLegend | #320008 |
| BV510 anti-mouse CD62L (MEL-14) | BD Biosciences | #563117 |
| BV605 anti-mouse Vα2 TCR (B20.1) | BD Biosciences | #747768 |
| PE anti-mouse CD132 (4G3) | BD Biosciences | #554457 |
| PE anti-mouse total Stat1 (N-terminus) (1/Stat1) | BD Biosciences | #558537 |
| AF647 anti-mouse Stat5 (pY694) (47/Stat5) | BD Biosciences | #612599 |
| PE anti-mouse Stat1 (pY701) (4a) | BD Biosciences | #612564 |
| BV605 anti-mouse CD69 (H1.2F3) | BD Biosciences | #563290 |
| APC anti-mouse CD62L (MEL-14) | ThermoFisher | #17-0621-82 |
| PE anti-mouse CD25 (PC61) | ThermoFisher | #12-0251-82 |
| FITC anti-mouse CD44 (5035-41.1D) | ThermoFisher | #MA1-81257 |
| PE-Cy7 anti-mouse CD127 (A7R34) | ThermoFisher | #25-1271-82 |
| FITC anti-mouse CD49d (R1-2) | ThermoFisher | #11-0492-82 |
| AF488 anti-mouse STAT5b (EPR16671) | Abcam | #ab199767 |
| *InVivo*MAb anti-mouse CD3ε (145-2C11) | BioXCell | BE0001-1 |
| *InVivo*MAb anti-mouse CD28 (37.51) | BioXCell | BE0015-1 |
| *InVivo*MAb anti-mouse IL-2 (S4B6-1) | BioXCell | BE0043-1 |
| **Chemicals and Reagents** | | |
| Paraformaldehyde 16% Aqueous Solution EM Grade | Electron Microscopy Sciences | #15710 |
| BD Phosflow™ Perm Buffer III | BD Biosciences | #558050 |
| FB-11: Fetal Bovine Serum, U.S. Source | Omega Scientific | #FB-11 |
| HyClone Phosphate Buffered Saline solution | Cytiva | #SH30256 |
| Percoll | Cytiva | #17089101 |
| RPMI 1640 Medium, no glutamine | ThermoFisher | #21870076 |
| Penicillin-Streptomycin (5,000 U/mL) | ThermoFisher | #15070063 |
| HEPES (1M) | ThermoFisher | #15630080 |
| Sodium Pyruvate (100 mM) | ThermoFisher | #11360070 |
| GlutaMAX™ Supplement | ThermoFisher | #35050061 |
| 2-Merceptoethanol | ThermoFisher | #21985023 |

| | | |
|---|---|---|
| Alexa Fluor™ 350 NHS Ester (Succinimidyl Ester) | ThermoFisher | #A10168 |
| CellTrace™ Violet Cell Proliferation Kit, for flow cytometry | ThermoFisher | #C34557 |
| PowerUp™ SYBR™ Green Master Mix for qPCR | ThermoFisher | #A25743 |
| Pierce™ DTT (dithiothreitol), No-Weight™ Format | ThermoFisher | #A39255 |
| NuPAGE™ Bis-Tris Mini Protein Gels, 4-12%, 1.0-1.5 mm | ThermoFisher | #NP0336 |
| NuPAGE™ MOPS SDS Running Buffer (20X) | ThermoFisher | #NP0001 |
| IL-2 Mouse Uncoated ELISA Kit with Plates | ThermoFisher | #88-7024-86 |
| Collagenase Type 4 | Worthington Biochemical Corporation | #LS004189 |
| Deoxyribonuclease I | Worthington Biochemical Corporation | #LS002139 |
| True-Nuclear™ Transcription Factor Buffer Set | BioLegend | #424401 |
| Recombinant Murine IL-2 | PeproTech | #212-12 |
| Recombinant Murine IL-15 | PeproTech | #210-15 |
| Recombinant Murine IFN-γ | PeproTech | #315-05 |
| OVA (257-264) Peptide Fragment | AnaSpec | #AS-60193-1 |
| OVA-Q4 Peptide, pQ4, SIIQFEKL, OVA (257-264) Variant | AnaSpec | #AS-64402 |
| EasySep™ Mouse CD8+ T Cell Isolation Kit | StemCell | #19853 |
| EasySep™ Mouse Naïve CD8+ T Cell Isolation Kit | StemCell | #19858 |
| EasySep™ Mouse Naïve CD4+ T Cell Isolation Kit | StemCell | #19765 |
| 4x Laemmli Sample Buffer | Bio-Rad Laboratories | #1610747 |
| Intercept (TBS) Blocking Buffers | LI-COR Biosciences | #927-60001 |
| Anti-SH2B3 antibody produced in rabbit | Sigma-Aldrich | #HPA005483 |
| Phospho-Stat5 (Tyr694) Antibody | Cell Signaling Technology | #9351S |
| β-Actin (8H10D10) Mouse mAb | Cell Signaling Technology | #3700S |
| RNeasy Mini Kit | Qiagen | #74104 |
| *Quick*-RNA Microprep Kit | Zymo Research | #R1050 |
| **Mouse Strains** | | |
| C57BL/6J (B6) | The Jackson Laboratory | #000664 |
| B6.Cg-Tg(Cd4-cre)1Cwi/BfluJ (CD4Cre) | The Jackson Laboratory | #022071 |
| B6.SJL-*Ptprc*$^{a}$*Pepc*$^{b}$/BoyJ (B6 CD45.1) | The Jackson Laboratory | #002014 |
| B6.Cg-*Gt(ROSA)*$^{26Sortm14(CAG-tdTomato)Hze}$/J (Ai14) | The Jackson Laboratory | #007914 |
| C57BL/6-Tg(TcraTcrb)1100Mjb/J (OT-I) | The Jackson Laboratory | #003831 |
| B6.Cg-Tg(TcraTcrb)425Cbn/J (OT-II) | The Jackson Laboratory | #004194 |
| C57BL/6-Tg(Ins2-TFRC/OVA)296Wehi/WehiJ (RIP-mOVA) | The Jackson Laboratory | #005431 |
| NOD.ShiLtJ (NOD) | The Jackson Laboratory | #001976 |

**Association and haplotype analysis.** Genomic DNA from T1D-affected sibling pairs (ASP) and trio families were obtained from the Type 1 Diabetes Genetics Consortium (T1DGC) as described previously[1]. Association analysis utilized data from T1DGC using genotypes for the 12q24 region. The Family-Based Association Test (FBAT) program (version 2.0.4) was used for single-marker association tests and haplotype analyses. Minor allele frequencies were estimated using PLINK (v.1.90).

**Mice.** All animal care and experimentation occurred with Institutional Animal Care and Use Committee approval at the SCRI Animal Facility. *Sh2b3*$^{-/-}$ and *Sh2b3*$^{fl/fl}$ mice were described previously[2]. All other strains were purchased from Jackson Laboratories (Bar Harbor, Maine): C57BL/6J (B6), B6.Cg-Tg(Cd4-cre)1Cwi/BfluJ (CD4Cre), B6.SJL-*Ptprc*$^{a}$ *Pepc*$^{b}$/BoyJ (B6 CD45.1), B6.Cg-*Gt(ROSA)26Sor*$^{tm14(CAG\text{-}tdTomato)Hze}$/J (Ai14), C57BL/6-Tg(TcraTcrb)1100Mjb/J (OT-I), B6.Cg-Tg(TcraTcrb)425Cbn/J (OT-II), C57BL/6-Tg(Ins2-TFRC/OVA)296Wehi/WehiJ (RIP-mOVA), and NOD/ShiLtJ (NOD). The NOD.*Sh2b3*$^{-/-}$ mice were generated by backcrossing *Sh2b3*$^{-/-}$ mice 15 generations to NOD mice. Genome wide SNP genotyping confirmed NOD sequences, including rs4225398 and rs3668978 flanking *Sh2b3* on chromosome 5. Intercrossing of heterozygous mice resulted in littermate control cohorts of *Sh2b3*$^{+/+}$, *Sh2b3*$^{+/-}$, and *Sh2b3*$^{-/-}$ NOD mice of each sex and genotype experimentally aged up to 40 weeks in diabetes incidence experiments. Mice were otherwise used at 6-12 weeks of age.

**Flow cytometry.** Antibody suspensions were made in PBS. When stained for surface markers, cells were resuspended in the fitting antibody suspension and incubated at 4C for 45 minutes, protected from the light. The targeted surface markers were CD4 (GDK1.5), CD8 (53-6.7), CD44 (IM7 and 5035-41.1D), CD62L (MEL-14), CD49d (R1-2), Vα2 TCR (B20.1), CD25 (PC61), CD69 (H1.2F3), CD122 (TM-β1), CD127 (A7R34), and CD132 (4G3). When stained for FOXP3 (150D), cells were fixed and permeabilized using the BioLegend True-Nuclear Transcription Factor Buffer Set. When stained for intracellular markers other than FOXP3, cells were fixed with 2% PFA at 37C for 10 minutes. The fixed cells were washed twice with FACS buffer (PBS + 2% FBS) before being permeabilized with BD Phosflow™ Perm Buffer III for 35 minutes at 4C. The cells were then washed twice with FACS buffer and resuspended in the fitting antibody suspension. The targeted intracellular markers were total STAT5 (EPR16671), total STAT1 (1/Stat1), phosphorylated STAT5 (pY694) (47/Stat5), and phosphorylated STAT1 (pY701) (4a). Prior to analysis, cells were washed free of the antibody suspension and resuspended in FACS buffer. Fluorescently labeled cells were acquired on BD LSR II or Fortessa (Becton Dickson) and analyzed using FlowJo (v9.7.6) (Treestar, Ashland, OR). Graphs and figures were prepared using GraphPad Prism v10.0.0 (Boston, MA) and Adobe Illustrator v16.0.0, respectively.

**Cell cultures.** Spleens, thymuses, and LNs were harvested from indicated mice and kept in complete RPMI media (RPMI-1640, 10% FBS, 1% penicillin-streptomycin, 1% HEPES, 1% sodium pyruvate, 1% GlutaMAX, and 0.1% 2-mercaptoethanol). Cells were isolated from the respective tissues using mechanical disruption and strained through a 40 µm filter. Spleen cells were treated with ACK lysis buffer for 5 minutes to lyse red blood cells before being resuspended in complete RPMI. In indicated experiments, total CD8+ T cells, naïve CD8+ T cells, or naïve $CD4^{+}$ T cells were isolated from spleen and/or LN cells with the respective StemCell isolation kit.

For cytokine stimulation, bulk splenocytes were plated at 1e6 cells/well in U-bottomed 96-well plates and stained for surface markers as described before being rested for 2 hours in 100µL/well of serum-free RPMI-1640. Suspensions of murine IL-2, IL-7[3], IL-15, and IFNgamma were prepared at a concentration of 50 ng/µL before being added 1:1 to the plated cells at the indicated time points for a final concentration of 25 ng/µL. For dose response, these concentrations were modified to suspensions of 50, 5, 0.5, and 0.05 ng/µL and final concentrations of 25, 2.5, 0.25, and 0.025 ng/µL. Following stimulation, the cells were fixed, permeabilized, and stained for intracellular pSTAT5 and pSTAT1 as described.

For plate-bound TCR stimulation, suspensions of anti-CD3 stimulatory antibody with or without anti-CD28 stimulatory antibody were made at indicated concentrations in PBS. In flat-bottomed 96-well plates, 50 µL of the indicated stimulatory antibody suspension was added to each well and the plate was coated overnight at 4C. After coating, excess suspension was removed and 100 µL of complete-RPMI was added to each well. Total *Sh2b3*$^{+/+}$ and *Sh2b3*$^{-/-}$ CD8+ T cells were plated at 2e6 cells/well. Naïve OT-I.*Sh2b3*$^{+/+}$ and OT-I.*Sh2b3*$^{-/-}$ CD8+ T cells were plated at 1e5 cells/well for flow cytometry analysis and at 2e5 cells/well for expression analysis. Naïve OT-II.*Sh2b3*$^{+/+}$ and OT-II.*Sh2b3*$^{-/-}$ CD4$^{+}$ T cells were plated at 5e4 cells/well for flow cytometry analysis. Cells plated for quantitative PCR expression analysis were separated into wells with anti-CD3 alone, with anti-CD3 and 10 µg/mL blocking anti-IL-2, and with 100 ng/mL murine IL-2 without anti-CD3. Cells plated for bulk RNA sequencing (RNAseq) were separated into wells with and without 10 µg/mL blocking anti-IL-2. Cells were cultured at 37C for the indicated length of time before being stained for surface markers as described or harvested for RNA isolation.

For antigen presenting cell (APC) TCR stimulation, splenocytes were isolated from B6 mice and irradiated at 3500rad to halt proliferation. Irradiated splenocytes were plated as feeder APCs at 5e6 cells/well in U-bottomed 96-well plates. SIINFEKL (N4) and the altered peptide ligand (APL) SIIQFEKL (Q4) were added to the feeder cells at concentrations from $10^{-10}$ to $10^{-7}$ M. Naïve OT-I.*Sh2b3*$^{+/+}$ and OT-I.*Sh2b3*$^{-/-}$ CD8+ T cells were loaded with CellTrace Violet according to manufacturer' protocol and plated with the feeder cells at 7.5e4 cells/well. Cells were cultured at 37C for 72 hours before being harvested and stained for surface markers as described. Upon harvest, the supernatant of the cultured cells was kept to assess IL-2 contents via an IL-2 mouse uncoated ELISA kit per manufacturer's protocol.

**Diabetes murine studies.** In adoptive transfer experiments, naïve OT-I CD8+ T cells were purified from spleen and cutaneous LN before being transferred retro-orbitally into sex-matched RIP-mOVA$^{+}$ hosts (7 – 14 weeks). When assessing proliferation of transferred cells, the donor cells were loaded with CellTrace Violet prior to transfer. Cells were transferred based on *Sh2b3* genotype, either in separate transfers in separate hosts or mixed 1:1 in a co-transfer into the same host. Cells were transferred at the indicated doses (7.5e5, 1.4e6, 1.5e6, 2e6, or 5e6 cells), suspended in 200 µL of chilled PBS. RIP-mOVA hosts were taken down after the indicated length of time (3, 5, 7, or 10 days) and tissues were harvested for flow cytometry or histology analysis. When processed for flow cytometry, spleens and LNs were processed as previously described. Pancreases were first mechanically disrupted by cutting into small pieces in pre-warmed digestion buffer (HBSS, 10% FBS, 2 mg/mL collagenase, and 10 µg/mL DNase) before being placed in an incubator shaker at 37C and 225 RPM for 35 minutes. The suspension was filtered through a 40 µm filter, with the resulting filtrate being pelleted and resuspended in 37% Percoll. The suspension was layered onto 70% Percoll and centrifuged for 25 minutes at 2600 RPM without brakes. Cells were isolated from the interphase of the density gradient before being washed in FACS buffer. Isolated cells were prepared for flow cytometry analysis as described. When processed for histology, harvested tissues were fixed in 10% neutral buffered formalin prior to Paraffin-embedding and sectioning with indicated staining and immunohistochemistry. Histology samples were independently assessed and graded for pattern of injury based on percent of total tissue injury, inflammation/necrosis, severity of involvement, and loss of insulin staining. Adoptive transfer and spontaneous NOD diabetes cohorts were monitored for hyperglycemia characterized by blood glucose > 300 mg/dL, with two consecutive readings requiring euthanasia per protocol. Blood glucose was measured in adoptive transfer recipients at 5, 7, 10, 14, and/or 28 days post-transfer. NOD cohorts’ urine was tested for glycosuria weekly starting at 8 weeks of age, then any glycosuria was confirmed with blood glucose measurement, repeated within 24 hours.

**Western blot analysis.** Protein was isolated from indicated cells by resuspending cell pellets in RIPA lysis buffer with added protease and phosphatase inhibitors. The resuspension was incubated on ice for 30 minutes before being centrifuged at 10,000 - 12,000 x g at 4C for 10 minutes before the supernatant was isolated. A solution of 4x Laemmli sample buffer and 200 µM DTT was made and added to the supernatant at 1:3 ratio, for a final concentration of 1x Laemmli buffer and 50 µM DTT. The solution was heated to 95C for 10 minutes before being cooled and loaded into a NuPAGE 4 to 12% Bis-Tris gel, ran in MOPS buffer at 150V for 90 minutes. The gel was transferred to a nitrocellulose membrane using the iBlot 2 system. The membrane was incubated in blocking buffer for 30 minutes at room temperature on a rocker. The blocking buffer was removed and the membrane was incubated in a suspension 1:1000 of rabbit anti-mouse SH2B3 in TBST overnight at 4C on a rocker. The membrane was washed with TBST five times before being incubated with a 1:10,000 suspension of anti-rabbit secondary for 1 hour at room temperature and washed five times with TBST. This process of primary antibody incubation, washing, secondary antibody incubation, and washing was repeated with rabbit anti-mouse pSTAT5 (Tyr694) and anti-mouse β-actin (with an anti-mouse secondary antibody used for the β-actin primary). The membrane was imaged using the Li-Cor Odyssey CLx.

**RNA-seq analysis**

RNA was isolated from indicated cells using the Qiagen RNeasy Kit or the Zymo *Quick*-RNA Microprep Kit per manufacturer’s protocol. For RT-PCR, cDNA was synthesized using the Thermo Scientific Maxima First Strand cDNA Synthesis Kit for RT-qPCR per manufacturer’s protocol. Murine *Sh2b3* and *Actb* transcripts were assessed by RT-PCR analysis using SYBR green master mix per manufacturer protocol. SH2B3 mRNA (ENSG00000111252) transcript sequence analysis was described previously[4,5]. Bulk mRNA-seq was performed via NovaSeq PE150 RNA sequencing and analyzed in the Biojupies cloud platform[6] or sequencing reads separately with data cleaning, differential expressed gene analysis and enrichment analysis as described[7].

*RNA-Seq data cleaning*. Sequences were quality-assessed with FastQC (v0.11.9)[8] and filtered with AdapterRemoval (v2.3.2)[9] to remove adapters and poor-quality sequences (score < 30, length < 15, ambiguous > 1). Sequences were aligned to the mouse genome (GRCm39 release 106) with STAR (v2.7.9a)[10] and quality-assessed with samtools flagstat (v1.7)[11] and Picard (v2.42.2)[11]. Alignments were filtered with samtools to remove PCR duplicates, unmapped, non-primary, and poor-quality alignments (MAPQ < 30). High-quality alignments were then quantified in gene exons using Subread featureCounts (v2.0.1)[12]. Data analysis and cleaning were done in the R version 4.2.2[13]. Libraries were filtered for sequence alignment of > 75%, median coefficient of variation of coverage < 0.9, and 2 standard deviations from the mean on principal components (PC) 1 and 2, thus resulting in 60 libraries for analysis. A batch effect was detected using principal component analysis (PCA) between experiments and was removed using CombatSeq from the R package sva[14]. Counts were normalized for RNA composition using the trimmed mean of M values normalization method and filtered to protein coding genes with at least 3 libraries containing at least 0.1 count per million (CPM). Finally, counts were converted to log2 CPM using R package voom[15].

*Differentially Expressed Gene (DEG) Analysis*. Experiment 1 and experiment 2 were analyzed separately. Pairwise contrasts were performed for each experiment to analyze contrasts between time/condition and genotype, including animal as a random effect (~ Time*Genotype + (1|Animal)) using the R package kimma[7]. Both experiments included a wild-type and knockout genotype. Experiment 1 contrasts for time included 0, 2, and 6 hours, and experiment 2 contrasts for time and condition included 0 and 24 hours with and without an anti-IL2 treatment. Contrasts were filtered for significance using an FDR < 0.25. Heatmaps using the R package ComplexHeatmap[16] were constructed for each experiment. For experiment 1, significant contrasts between wild-type and knockout within time points 2 or 6 hours as well as significant interaction term genes (Time:Genotype) were selected (FDR < 0.25). Log2 CPM were root-mean-squared scaled and complete linkage hierarchically clustered. For experiment 2, significant contrasts between wild-type and knockout within 24 hours as well as significant interaction term genes (Time:Genotype) were selected (FDR < 0.25). Log2 CPM were root-mean-squared scaled and complete linkage hierarchically clustered.

*DEG Enrichment Analysis*. DEGs per contrast were extracted from experiment 1 and experiment 2 to perform enrichment against the Human Molecular Signatures Database (MSigDB 2023.1.Hs) including Gene Ontology (GO) C5 biological pathways and Hallmark pathways using Fisher’s exact test in BIGprofiler from the R package SEARchways[15]. Significant pathways were filtered for using an FDR < 0.25 and > 5 DEGs present per pathway.

**Stastical analysis.** All statistical analysis used GraphPad Prism version 7.0b except where noted. All specific statistical tests and *P*-values are indicated in the relevant figures.
